# Manipulating rice canonical Gα and extra-large G protein subunits for improved agronomic traits

**DOI:** 10.1101/2024.08.17.608385

**Authors:** Christian F. Cantos, Sarah M. Assmann

## Abstract

Rice productivity is fundamentally linked to its architecture, governed by signaling networks including those based on heterotrimeric G proteins. In this study, we investigated the individual gene impacts and genetic interactions of the canonical Gα gene (*RGA1*), and the non-canonical extra-large Gα genes (*OsXLG1, OsXLG3a, OsXLG3b, OsXLG4*) in controlling plant architecture. We generated *OsXLG* mutants using CRISPR/Cpf1 gene editing in Nipponbare (WT) and *d1*, a Nipponbare null mutant of *RGA1*. We then phenotyped 25 different genotypes in the greenhouse for 19 different agronomic traits. In wild type (WT), mutations in *RGA1*, *OsXLG3a*, *OsXLG3b*, or *OsXLG4*, as well as any combination of Gα genes, resulted in a shorter stature, a desirable trait. Mutations in *OsXLG1* and *OsXLG4* increased the number of spikelets and grains per panicle, showcasing advantageous traits that led to higher yield. Mutations in *OsXLG3a*, *OsXLG3b*, any combination of *OsXLGs*, or any *OsXLG* combined with the *d1* mutation, reduced seed production and yield. Flag leaf width was the only trait influenced solely by RGA1. *RGA1* transcript abundance in the *osxlg* mutants was positively correlated with height, culm length, panicle exsertion, and harvest index, implicating OsXLG regulation of *RGA1* expression as an underlying mechanism. Overall, increased *RGA1* expression is correlated with more favorable reproductive traits but less favorable vegetative traits. Our study reveals the complex interaction of RGA1 and OsXLGs within the signaling networks that shape rice architecture, from vegetative to post-harvest stages. Our results suggest modulation of *RGA1, OsXLG1, OsXLG3a,* or *OsXLG4* expression as strategies to enhance yield.

## Introduction

Rice (*Oryza sativa*) is a staple food crop that sustains a significant portion of the world’s population. The demand for rice production is steadily rising due to increasing human population and drastic changes in the environment. Optimizing plant architecture is crucial for increasing rice productivity and acclimation to diverse environments (Ćalić *et al*., 2022; Liu *et al*., 2018).

Heterotrimeric G proteins, G proteins hereafter, are composed of Gα, Gβ, and Gγ subunits and are key components in cellular signaling (Assmann, 2005; Cantos *et al*., 2023). In the canonical paradigm, G protein signaling begins with the activation of G protein-coupled receptors (GPCRs) by extracellular ligands. Upon ligand binding, GPCR-mediated activation triggers the exchange of GDP (guanosine diphosphate) for GTP (guanosine triphosphate) bound to the Gα subunit, leading to the dissociation of Gα-GTP from the Gβγ dimer. The transition between the inactive Gα-GDP-βγ and active Gα-GTP and Gβγ states serves as a molecular switch, modulating downstream signaling cascades through interactions with effector proteins that regulate diverse processes (Papasergi-Scott *et al*., 2024).

In contrast to the extensive variety of G protein subunits and GPCRs found in mammals (Perfus-Barbeoch *et al*., 2004), plants possess a more streamlined set of G protein components, featuring plant-specific subunits such as the extra-large Gα (XLG) and unconventional Gγ proteins, and with Receptor-Like Kinases (RLKs) likely serving as major functional analogs to GPCRs in signal transduction (Aranda-Sicilia *et al*., 2015; Maruta *et al*., 2021). Plants also display distinct G protein biochemistry; for example, while the Arabidopsis Gα subunit GPA1 shares structural parallels with mammalian Gα proteins, it exhibits rapid activation due to accelerated nucleotide exchange (Gookin *et al*., 2023).

Research in the model plant species Arabidopsis, maize and rice has revealed the influence of G proteins on numerous aspects of plant development, including plant architecture. In Arabidopsis, the canonical Gα protein, GPA1, and non-canonical extra-large G-proteins (AtXLGs), play important roles in development and stress responses. Null mutants of *gpa1* exhibit notable phenotypic changes, including shorter and wider hypocotyls, flowers, and siliques, alongside a decreased stomatal density (Ge *et al*., 2008; Ullah *et al*., 2002; Zhang *et al*., 2008). Similarly, *AtXLG2* and *AtXLG3* T-DNA knockout mutants and an *AtXLG1* T-DNA knockdown mutant implicate AtXLGs in suppressing primary root growth (Ding *et al*., 2008), enhancing root waving (Pandey *et al*., 2008), and inhibiting stomatal proliferation (Chakravorty *et al*., 2015). Moreover, physical interaction between AtXLGs and PUB family E3 ubiquitin ligases adds complexity to their roles in hormonal regulation and developmental processes (Wang *et al*., 2017). Higher-order mutants of *AtXLGs* and interacting *PUBs* display a reduced sensitivity to cytokinin, manifest decreased stamen number and exhibit defects in tapetum development resulting in lower fertility (Wang *et al*., 2017).

In maize, the canonical Gα protein, CT2, and non-canonical ZmXLGs show both overlapping and independent roles in regulating plant height and meristem development (Wu *et al*., 2018). The *ct2* mutant displays short stature with an enlarged shoot apical meristem (SAM) and fasciated ears, while *zmxlg* triple mutants exhibit seedling lethality, confirming the importance of XLGs for growth and viability (Wu *et al*., 2018). Interestingly, the reduced height and enlarged SAM phenotype in *ct2* are enhanced when combined with any pair of *zmxlgs*, suggesting overlapping roles of ZmXLGs and CT2 in controlling plant height and cell proliferation in the SAM (Wu *et al*., 2018). However, knocking out any pair of *zmxlgs* does not enhance ear fasciation in the *ct2* mutant background, indicating an independent role of CT2. Rice has one canonical Gα gene (*RGA1*) and four non-canonical *OsXLG* genes (*OsXLG1, OsXLG3a, OsXLG3b,* and *OsXLG4*) (Cantos *et al*., 2023). The canonical *RGA1* gene has been extensively studied, revealing critical roles in plant architecture and various abiotic and biotic stress responses (Cantos *et al*., 2023). Several historically isolated null mutants of *RGA1*, known as *d1*, show shorter stature with broad and erect leaves, compact panicles, and small rounded seeds (Ashikari *et al*., 1999; Oki *et al*., 2009). Additionally, CRISPR mutants of *RGA1* show similar phenotypes as the *d1* mutant as well as early heading and seed setting, with reduced grain weight (Cui *et al*., 2020). The short stature and broad, erect leaf phenotypes are advantageous traits that can resist lodging (Ferrero-Serrano *et al*., 2019), and promote light harvesting for photosynthesis (Ferrero-Serrano *et al*., 2018), respectively, which could contribute favorably to harvest index and yield (Ferrero-Serrano *et al*., 2019). However, compact panicles could negatively affect pollination and grain formation, resulting in an overall yield reduction (Parida *et al*., 2022). *d1* plants are drought-resistant with increased photoprotection and decreased photoinhibition (Ferrero-Serrano *et al*., 2018), and increased mesophyll conductance (Zait *et al*., 2021), which contribute to increasing photosynthesis and water-use efficiency (Ferrero-Serrano and Assmann, 2016; Yantong *et al*., 2022; Zait *et al*., 2021).

Mutational analyses indicate that OsXLGs also regulate agronomically important traits such as height, tiller number, panicle number, grains per panicle, and grain size (Biswal *et al*., 2021; Cui *et al*., 2020; Zhao *et al*., 2022). OsXLGs also contribute to tolerance against chilling (Cui *et al*., 2020), salinity (Biswal *et al*., 2021; Cui *et al*., 2020), and drought stress (Cui *et al*., 2020). Moreover, OsXLGs also play a role in disease resistance against bacterial leaf blight (Biswal *et al*., 2021; Zhao *et al*., 2022) and rice blast (Zhao *et al*., 2022). While roles of RGA1 and OsXLGs in rice architecture have been individually dissected, their functional partitioning and overlapping roles as well as possible coordinated expression between these Gα proteins has not been studied.

We investigated the roles of all five Gα genes and their potential genetic interactions by generating CRISPR mutants of *OsXLGs* in two Nipponbare cultivars: Nipponbare wild type (with functional RGA1) and the *d1* mutant (with nonfunctional RGA1). In this *d1* mutant, a single base pair substitution (G^508^ to A^508^) in exon 7 of *RGA1* gene introduces a premature stop codon, and immunoblot assays confirm the absence of RGA1 protein (Ashikari *et al*., 1999; Oki *et al*., 2009). We assessed the phenotypic effects of *OsXLG* single, double, and triple mutants, as well as the combination of editing each *OsXLG* mutation in the *d1* background. This research contributes to our understanding of G protein signaling in plants and provides insights into potential strategies for optimizing crop architecture to enhance yields.

## Materials and Methods

### Generation of CRISPR lines

We used the CRISPR/Cpf1 gene editing system to generate *OsXLGs* mutants in Nipponbare wild type (WT) and *d1* mutant backgrounds by employing two approaches: 1) a single CRISPR RNA (crRNA) target per *OsXLG,* 2) multiple crRNA targets per *OsXLG* (SI, Figure 1a). We identified crRNA target sites using the Benchling CRISPR design tool (https://www.benchling.com/crispr) and selected three crRNA sites per *OsXLG* with no off-targets based on NCBI BLAST off-target analysis (https://blast.ncbi.nlm.nih.gov/Blast.cgi) (SI, Table 1).

**Figure 1.**
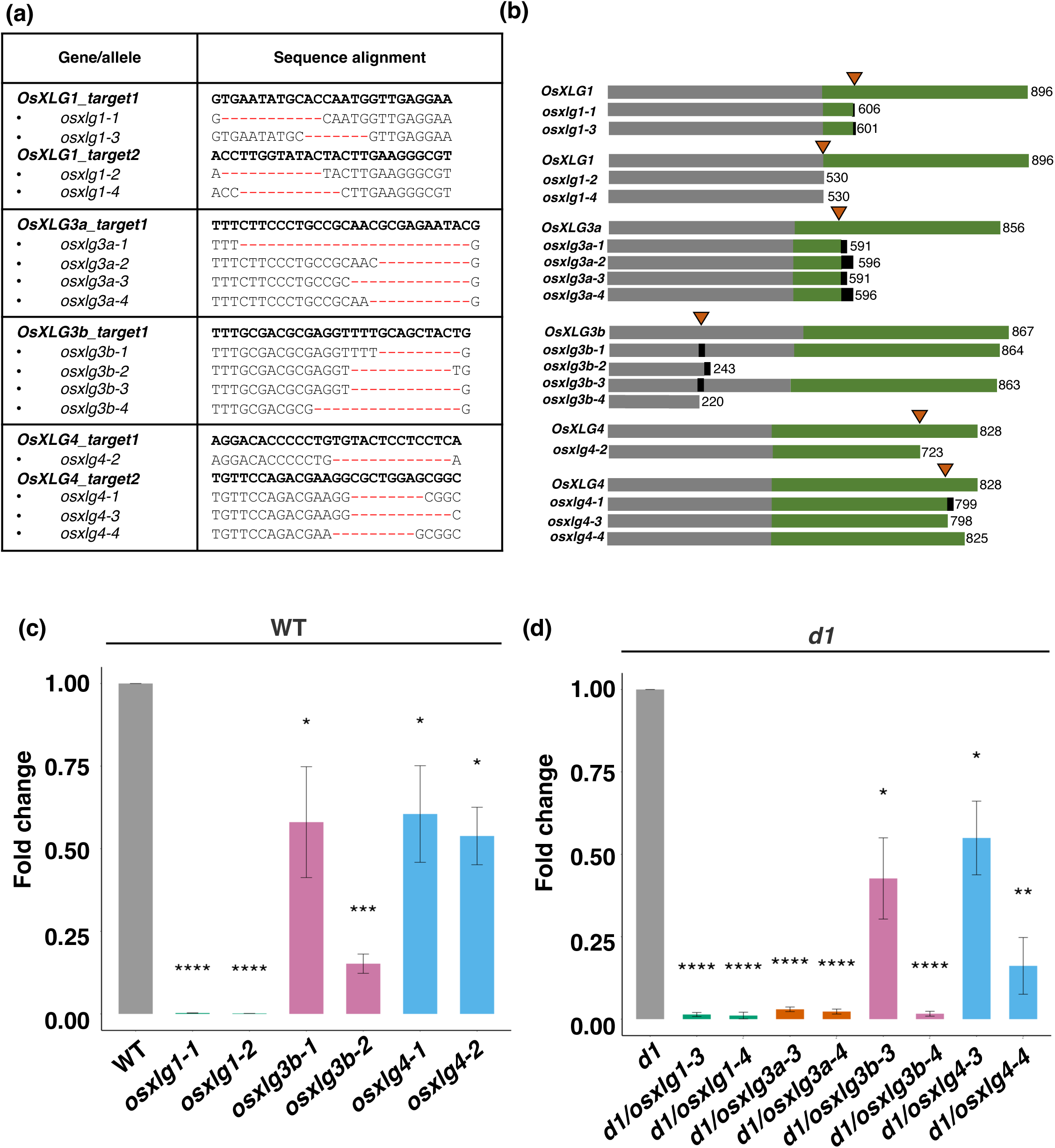
Schematic representation, sequences, and gene expression analysis of extra-large G protein (*OsXLG*) mutant alleles generated by CRISPR/Cpf1 gene editing. Alleles −1 and −2 are WT background; alleles −3 and −4 are *d1* mutant background. (a) Sequences of the *osxlg* mutants generated by CRISPR/Cpf1 gene editing. (b) Schematic representation of the OsXLG proteins and the predicted truncations generated by CRISPR/Cpf1 gene editing. Orange arrowheads indicate the position of the CRISPR RNA target sites and numbers indicate the lengths of the corresponding amino acid sequences. Grey bars-N-terminal domain; Green bars-Gα domain; Black bars-position of predicted frameshift mutations. (c-d) Relative expression levels (fold change) of CRISPR mutants determined by quantitative reverse transcription polymerase chain reaction (qRT-PCR), as compared to background controls (WT or *d1*) Data are presented as mean values ± standard error of the mean from three technical replicates and three biological replicates. Statistical significance is indicated as follows: *, P < 0.05; **, P < 0.01; ***, P < 0.001; ****, P < 0.0001. p-values were generated using Student’s t-tests to assess the statistical significance of the observed differences between the mutant and control groups.

For the single crRNA target per *OsXLG* approach, we used plasmids from Addgene: pYPQ141 (#86197, crRNA and ribozyme expression array), pYPQ230 (#86210, LbCpF1 nuclease vector) and pYPQ203 (#86207, T-DNA vector) (SI, Figure 1b). crRNA_target1 (SI, Table 1) was selected for each *OsXLG* and concatenated with direct repeat sequence (DR) with BamHI enzyme overhang and introduced to pYPQ141 by restriction enzyme digestion-ligation. The pYPQ141_crRNA_target1 vector was introduced to pYP230, followed by integration into pYPQ203 through Gateway cloning using the LR Clonase system (ThermoFisher). For the multiple crRNA targets per *OsXLG* approach, we used the single transcriptional unit (SSTU) system of CRISPR/Cpf1 designed for multiplex editing (Wang *et al*., 2018) (SI, Figure 1c). Three crRNA target sites per *OsXLG* were concatenated with the DR repeats with BamHI enzyme overhangs. The concatenated sequences were introduced to pSSTU_LbCpF1 vectors by restriction enzyme digestion-ligation. The CRISPR constructs were sequence-confirmed and then introduced into WT and *d1* mutant rice calli through *Agrobacterium*-mediated stable rice transformation (Nishimura *et al*., 2006). The transformations with the CRISPR constructs for the single target approach for all OsXLGs in both WT and *d1* backgrounds, as well as transformations with the CRISPR constructs for the multiple targets approach against *OsXLG3b* and *OsXLG1* genes in WT, were performed in our laboratory. The transformation of CRISPR constructs for the multiple targets approach for *OsXLG3a* and *OsXLG4* were conducted at Cornell’s transformation facility.

### Mutant screening and T-DNA analysis

Genomic DNA was extracted from T0 plants using a modified CTAB protocol. The *OsXLG* crRNA target sequences were amplified using gene-specific primers (SI, Table 2) and Q5 Hi-Fi DNA polymerase (NEB). The PCR amplicons were subjected to Sanger sequencing and analyzed using CRISP-ID online software (Dehairs *et al*., 2016) to identify mutations. T0 plants with desired mutations were maintained to the T1 generation. In the T1 generation, homozygous and CRISPR-free (T-DNA PCR negative) mutants were identified and maintained for production of T2 generation seeds (SI, Table 3). CRISPR-free mutants were identified by determining the presence or absence of T-DNA in the mutant’s genome by PCR amplification of the *hygromycin phosphotransferase* (*hpt*) gene within the T-DNA (Table 1 and SI, Table 3).

**Table1.** *OsXLG* naming system for Biswal et al. (2021) *osxlg* mutants.

| Biswal et al. (2021) CRISPR mutants | *OsXLG* name Cantos et al. (2024) | Allele designation in this study | Number of CRISPR free plants | Note |
| --- | --- | --- | --- | --- |
| *osxlg4-1* | *OsXLG1* | *osxlg1-5* | 0/5 | SI, Figure 3 |
| *osxlg2-1* | *OsXLG3a* | *osxlg3a-5* | 5/5 | Included in the discussion |
| *osxlg1-1* | *OsXLG3b* | *osxlg3b-5* | 0/5 | SI, Figure 3 |
| *osxlg2-5, 4-2)*<br>*osxlg1-2, 4-2*<br>*osxlg1-3, 4-3* | *OsXLG1/OsXLG3a*<br>*OsXLG1/OsXLG3b*<br>*OsXLG1/OsXLG3b* | *osxlg1-6/3a-6*<br>*osxlg1-6/3b-6*<br>*osxlg1-7/3b-7* | 5/5<br>5/5<br>5/5 | SI, Figure 3 |
| *osxlg1,2,4-3*<br>*osxlg1,2,4-5*<br>*osxlg1,2,4-6* | *OsXLG1/3a/3b*<br>*OsXLG1/3a/3b*<br>*OsXLG1/3a/3b* | *osxlg1/3a/3b* | 1/5<br>3/5<br>4/5 | CRISPR-free lines are consolidated for discussion as *osxlg1/3a/3b* |

### Plant materials and greenhouse conditions

Rice seeds of twenty-five genotypes, consisting of the T2 generation of single, double, triple, *osxlg* and *d1/osxlg* mutants, as well as WT and *d1* mutants, were germinated in petri dishes on moist filter paper and maintained in a growth chamber (Percival, USA) under controlled conditions (30°C, 16/8h light/dark cycle). After two weeks, the seedlings were transplanted into 2.6L pots filled with a 1:1 ratio of soil (C/V Mix, Jolly Gardener PRO-LINE, USA) and granular slow-release fertilizer (Osmocote plus). Plants were grown in the Penn State Agricultural Science greenhouse from March to November of 2022 with temperatures ranging from 26-32°C and humidity levels at 60-70%. Iron chelate supplement (Sequestrene 330) was applied weekly for the first month, and 20-10-20 (nitrogen–phosphorus–potassium) fertilizer (Jack’s Peat Lite Special) was applied weekly throughout the vegetative stage.

### Phenotype measurement

For phenotyping, five replicates of two-week-old seedlings of each mutant and their corresponding cultivar background controls (WT, *d1*) were measured, for a total of 125 plants. Vegetative phenotypes were measured for each replicate during the 8th week prior to flowering: plant height, tiller number, flag leaf length and width, and culm length. Days to heading was recorded and all plants were grown to maturity. Mature panicles were harvested, and reproductive phenotypes were recorded: panicle number, panicle length (exsertion and axis length), spikelets per panicle, grains per panicle, and spikelet fertility (total number of filled grains/ total number of spikelets). Seeds were collected and measured for 100-grain weight and total grain weight per plant (yield). Grain size analysis was conducted on twenty-five seeds per genotype using PTRAP software (A L-Tam *et al*., 2013). For shoot biomass, the entire above-ground shoot material was harvested after panicle collection, dried for one week in a 65 ^o^C oven, and weighed. Harvest index (HI) was calculated using the formula: HI = Yield (g)/ Dry shoot biomass (g).

### RNA isolation and qRT-PCR

Three biological replicates of entire non-flag leaves from 4-week-old rice plants of each genotype were harvested. Total RNA was extracted using the TRIzol RNA extraction kit (ThermoFisher). cDNA was synthesized using the SuperScript III First-Strand Synthesis system (ThermoFisher). Quantitative RT-PCR was performed on an iQ5 Real-Time system (Bio-Rad). Primer pairs downstream of each CRISPR *OsXLG* target site were used; primer pairs for *RGA1* were against the 3’ region of the CDS. Primers were validated using WT leaf samples to confirm their functionality. The relative expression level of each Gα gene was normalized using the housekeeping gene *GAPDH*. The fold change (log_2_ change of the ^ΔΔ^Ct) was calculated to determine the relative changes in Gα gene expression. All primer sequences are provided in SI, Table 2.

### Statistical analysis

Statistical analysis of significant differences between genetic backgrounds and *osxlg* mutants was conducted using Student’s t-tests in the R statistical programming language (www.R-project.orgs). Linear regression analysis was conducted using the “lm” function in R to assess the relationship between phenotypic traits and Gα gene expression. Error bars represent standard errors, and p-values are denoted as follows: *, p < 0.05; **, p < 0.01; ***, p < 0.001, ****, p < 0.0001.

### Evaluation of OsXLG-RGA1 interactions

To study how *OsXLGs* and *RGA1* mutations interact phenotypically, we initially averaged the phenotype values for the alleles of each *OsXLG* gene. After establishing these averages, we calculated the effects of each *osxlg* and the *d1* mutation by comparing these values to those for the Nipponbare wild type. Then, we calculated the expected combined mutational effect, assuming independence, by adding the individual mutational effects. We compared this value with the actual values seen in plants with both mutations, i.e., the *osxlg* double mutants in the Nipponbare background, and the *osxlg* single mutants in the *d1* background. Based on these comparisons, we determined whether the interaction was synergistic (greater than the additive effect), additive (equal to the calculated effect), partially redundant (less than the additive effect but still significant), and whether there were suppressive, enhancing, or antagonistic effects of Gα mutations.

## Results

### Generation of novel alleles of *OsXLGs*

Independent CRISPR edited lines were generated for *OsXLG1*, *OsXLG3a*, *OsXLG3b* and *OsXLG4* in both Nipponbare (WT) and *d1* mutant backgrounds (SI, Table 3). CRISPR mutations were detected at each of the *OsXLG’s* crRNA target sites with mutation efficiencies ranging from 35.7-100% (SI, Table 3). The mutations identified were mostly deletions (Figure 1a and SI, Table 4) that produced either in-frame deletions of several amino acids or frameshift mutations resulting in premature stop codons predicted to truncate the OsXLG protein (Figure 1b). For each *OsXLG* gene, we selected two independent lines to investigate for agronomic analyses, following the criterion of producing two distinct predicted OsXLG proteins (Figure 1b and SI, Table 4). Homozygous and CRISPR-free mutants were selected from the T1 generation and propagated to produce T2 generation plants, which were evaluated in the greenhouse experiment (SI, Table 3). In our nomenclature, Allele-1 and −2 are in WT background, while Allele-3 and −4 are in *d1* mutant background. The *OsXLG* alleles of CRISPR lines in WT background are *osxlg1-1* (11 bp deletion) and *osxlg1-2* (11 bp deletion), *osxlg3a-1* (25 bp deletion) and *osxlg3a-2* (10 bp deletion), *osxlg3b-1* (9 bp deletion) and *osxlg3b-2* (11 bp deletion), *osxlg4-1* (8 bp deletion) and *osxlg4-2* (13 bp deletion). The *OsXLG* alleles of CRISPR lines in the *d1* mutant background are *osxlg1-3* (7 bp deletion) and *osxlg1-4* (11 bp deletion), *osxlg3a-3* (13 bp deletion) and *osxlg3a-4* (11 bp deletion), *osxlg3b-3* (12 bp deletion) and *osxlg3b-4* (16 bp deletion), *osxlg4-3* (11 bp deletion) and *osxlg4-4* (9 bp deletion) (Figure 1a and SI, Table 4). Interestingly, none of the lines we generated of *OsXLG3a* in the WT background survived past the T0 generation (SI, Figure 2, and SI, Table 3).

**Figure 2.**
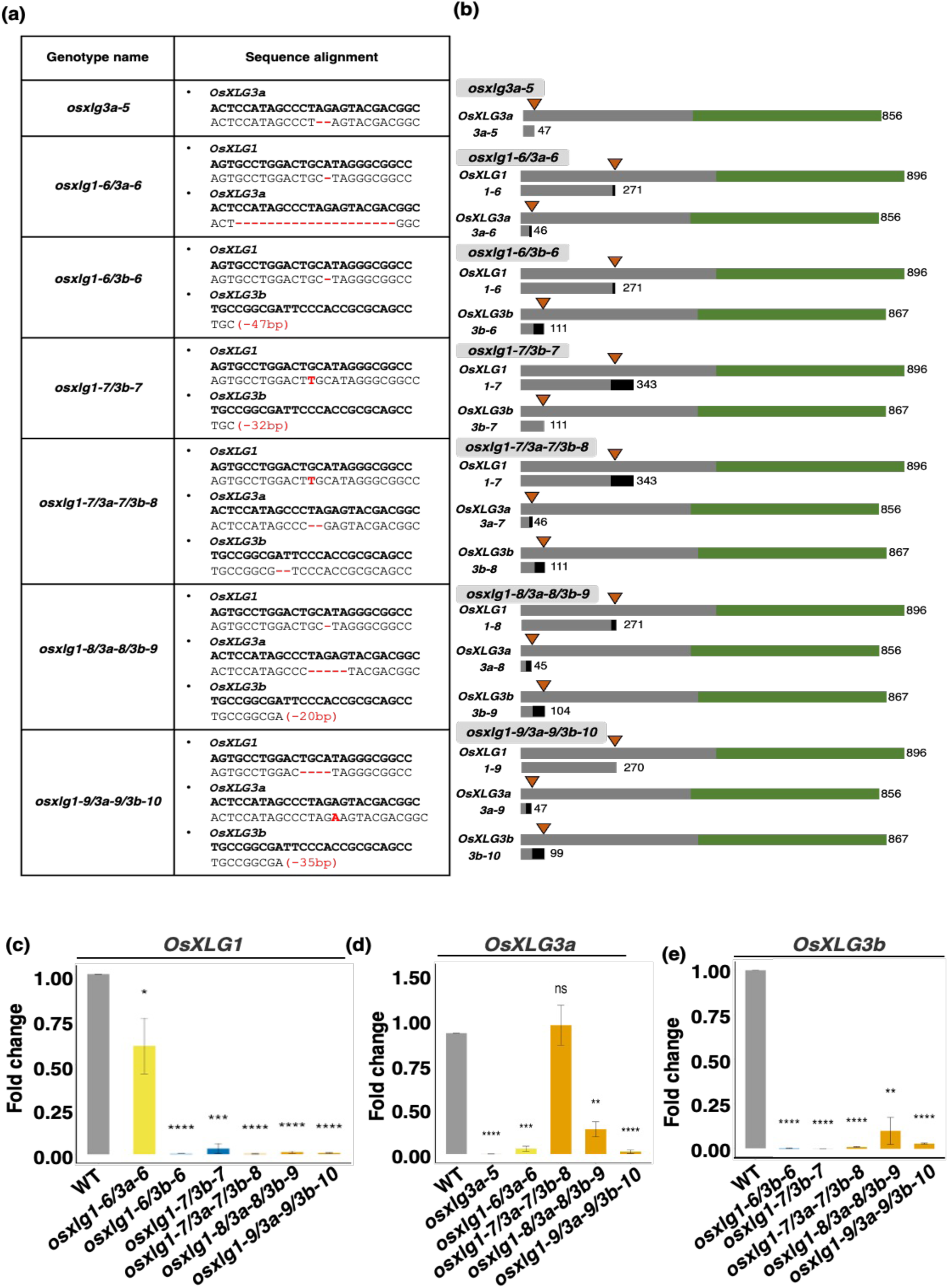
Schematic representation, sequences, and gene expression analysis of extra-large G protein (*OsXLG*) mutants from Biswal, et al. (2021) All of these CRISPR *osxlg* alleles are in the WT background. (a) Sequences of the CRISPR *osxlg* alleles. (b) Schematic representation of the OsXLG proteins and the predicted proteins of each CRISPR *osxlg* allele. Orange arrowheads indicate the position of the CRISPR RNA target sites and numbers indicate the lengths of the corresponding amino acid sequences. Grey bars-N-terminal domain; Green bars-Gα domain; Black bars-position of predicted frameshift mutations. (c-e) Relative expression levels (fold change) of CRISPR mutants determined by quantitative reverse transcription polymerase chain reaction (qRT-PCR), as compared to control (unedited) plants. Data are presented as mean values ± standard error of the mean from three technical replicates and three biological replicates. Statistical significance is indicated as follows: ns, nonsignificant; *, P < 0.05; **, P < 0.01; ***, P < 0.001; ****, P < 0.0001. p-values were generated using Student’s t-tests to assess the statistical significance of the observed differences between the mutant and control groups.

We also included in our experiments some of the CRISPR *osxlg* mutants from the study by (Biswal *et al*., 2021). We use our published, phylogenetically-based *OsXLG* naming system (Cantos *et al*., 2023)to assign names to these CRISPR mutants: *osxlg1-5* for their *osxlg4-1*, *osxlg3a-5* for their *osxlg2-1*, *osxlg3b-5* for their *osxlg1-1*, *osxlg1-6/3a-6* for their *osxlg2-5,4-2, osxlg1-6/3b-6* and *osxlg1-7/3b-7* for their *osxlg1-2, 4-2* and *osxlg1-3,4-3, osxlg1-7/3a-7/3b-8* for their *osxlg1,2,4-3*, *osxlg1-8/3a-8/3b-9* for their *osxlg1,2,4-5,* and *osxlg1-9/3a-9/3b-10* for their *osxlg1,2,4-6* mutants (Table 1). Figure 2a and 2b shows the mutation type and predicted protein of each *OsXLG* allele of the mutants from Biswal et al. (2021) that we analyzed. We also determined whether the CRISPR mutants sourced from Biswal et al. (2021) were CRISPR-free (Table 1). We discovered that mutant plants such as *osxlg3a-5*, *osxlg1-6/3a-6*, *osxlg1-6/3b-6*, and *osxlg1-7/3b-7* lack CRISPR elements. However, in the mutants *osxlg3b-5* and *osxlg1-5*, CRISPR elements were identified in 4 of 5 and 3 of 5 replicates, respectively (Table 1). Regarding the *osxlg1-7/3a-7/3b-8, osxlg1-8/3a-8/3b-9,* and *osxlg1-9/3a-9/3b-10* mutants, CRISPR elements were present in 4 of 5, 2 of 5, and 1 of 5 replicates, respectively. In Figures 2-7, we present results only from the CRISPR-free lines; our results from the non-CRISPR-free lines of Biswal et al. (2021) are presented in SI, Figure 3. Additionally, we consolidated data from all CRISPR-free lines of the triple mutant genotypes, which we designated as *osxlg1/3a/3b*, for discussion since the predicted impacts of the edits on protein structures are highly similar.

**Figure 3.**
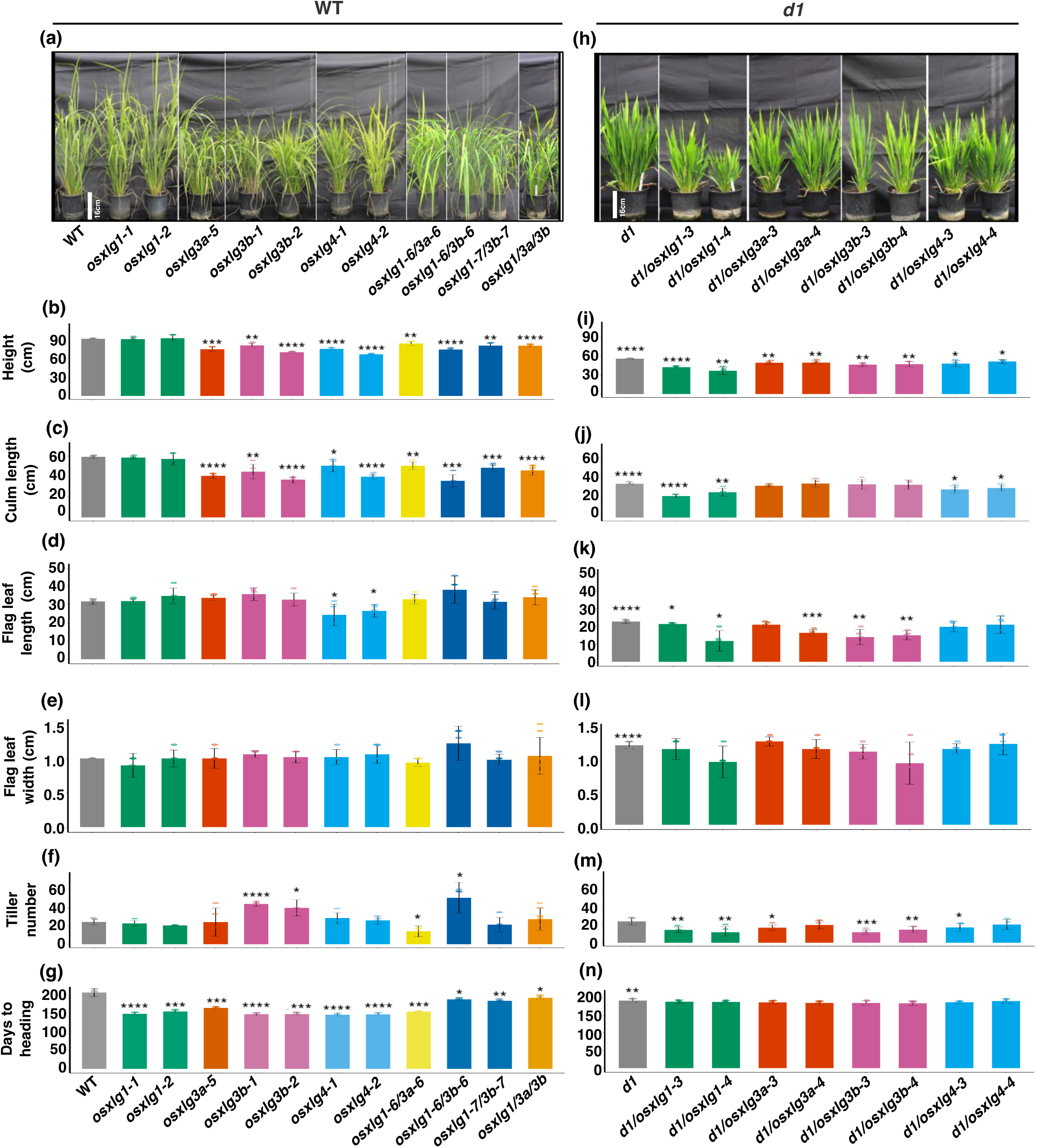
Effects of *OsXLG* mutations on vegetative phenotypes. (a, h) Images of the vegetative plant architecture of the *OsXLG* mutants. (b-g) Vegetative traits of *OsXLG* mutants in WT background. (i-n) Vegetative traits of *OsXLG* mutants in *d1* mutant background. The *d1* mutant significance level is indicated as compared to WT; significance levels for all other mutants are relative to their genotypic background. Statistical significance levels are indicated as follows: *, P < 0.05; **, P < 0.01; ***, P < 0.001, ****, P < 0.0001.

**Figure 4.**
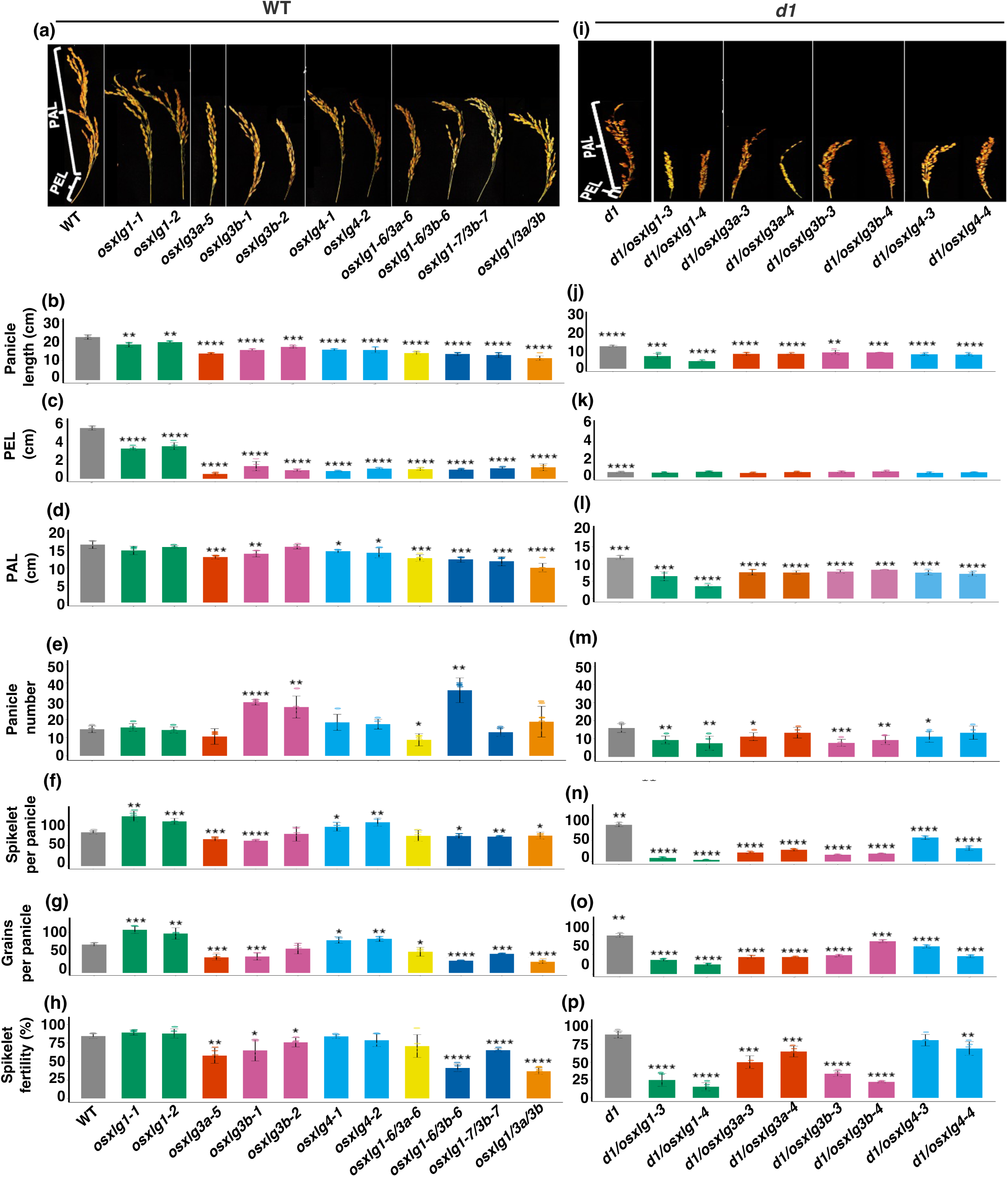
Effects of *OsXLG* mutations on reproductive phenotypes. (a, i) Images of panicle morphology of the *OsXLG* mutants. (b-h) Reproductive traits of *OsXLG* mutants in WT background. (j-p) Reproductive traits of *OsXLG* mutants in *d1* mutant background. The *d1* mutant significance level is indicated as compared to WT; significance levels for all other mutants are relative to their genotypic background. Statistical significance levels are indicated as follows: *, P < 0.05; **, P < 0.01; ***, P < 0.001, ****, P < 0.0001.

**Figure 5.**
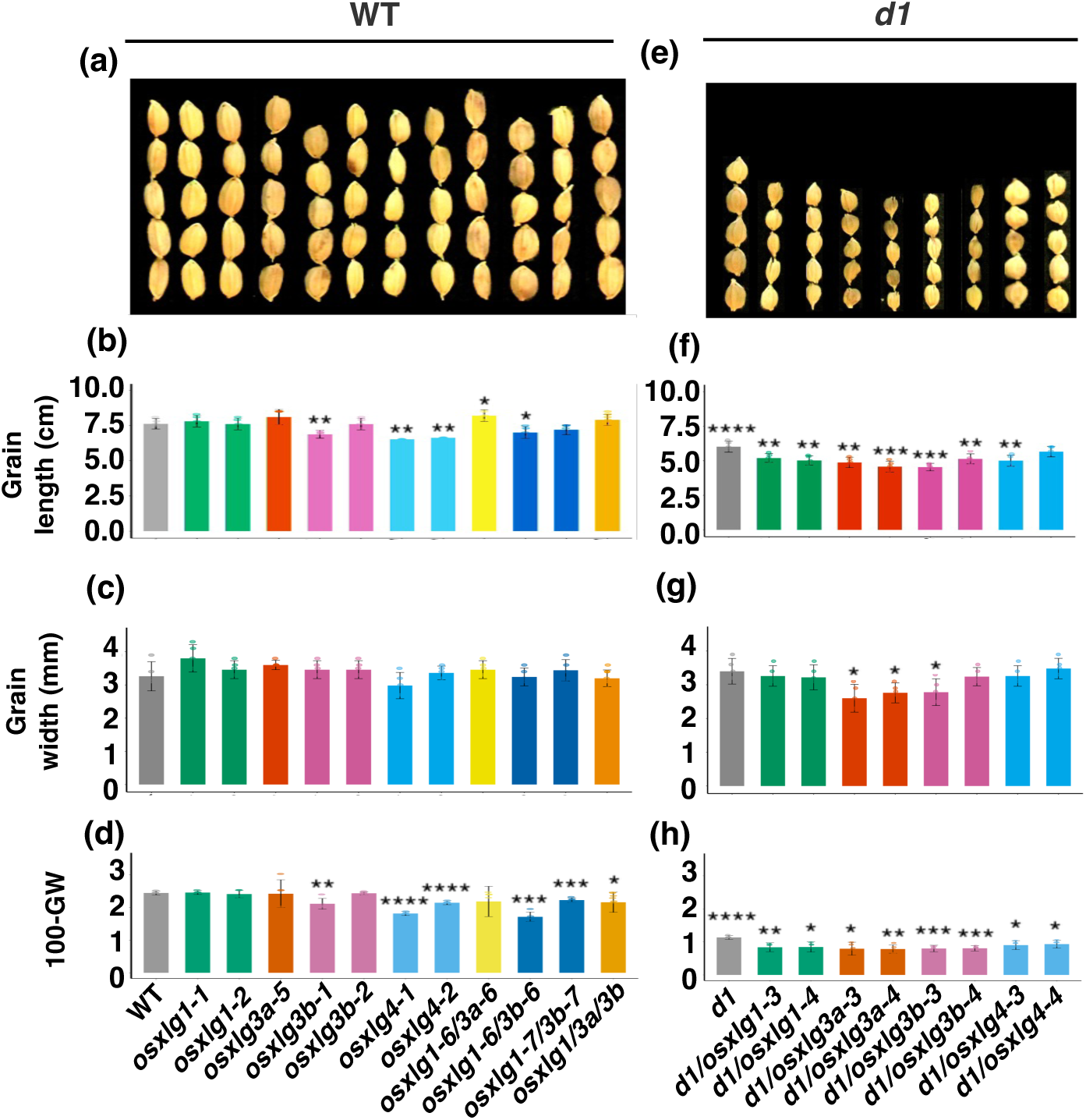
Effects of *OsXLG* mutations on grain phenotypes. Images and quantitation of grain length, grain width and grain weight of the *OsXLG* mutants in WT (a-d) and *d1* mutant (e-f) backgrounds. The *d1* mutant significance level is indicated as compared to WT; significance levels for all other mutants are relative to their genotypic background. Statistical significance levels are indicated as follows: *, P < 0.05; **, P < 0.01; ***, P < 0.001, ****, P < 0.0001.

**Figure 6.**
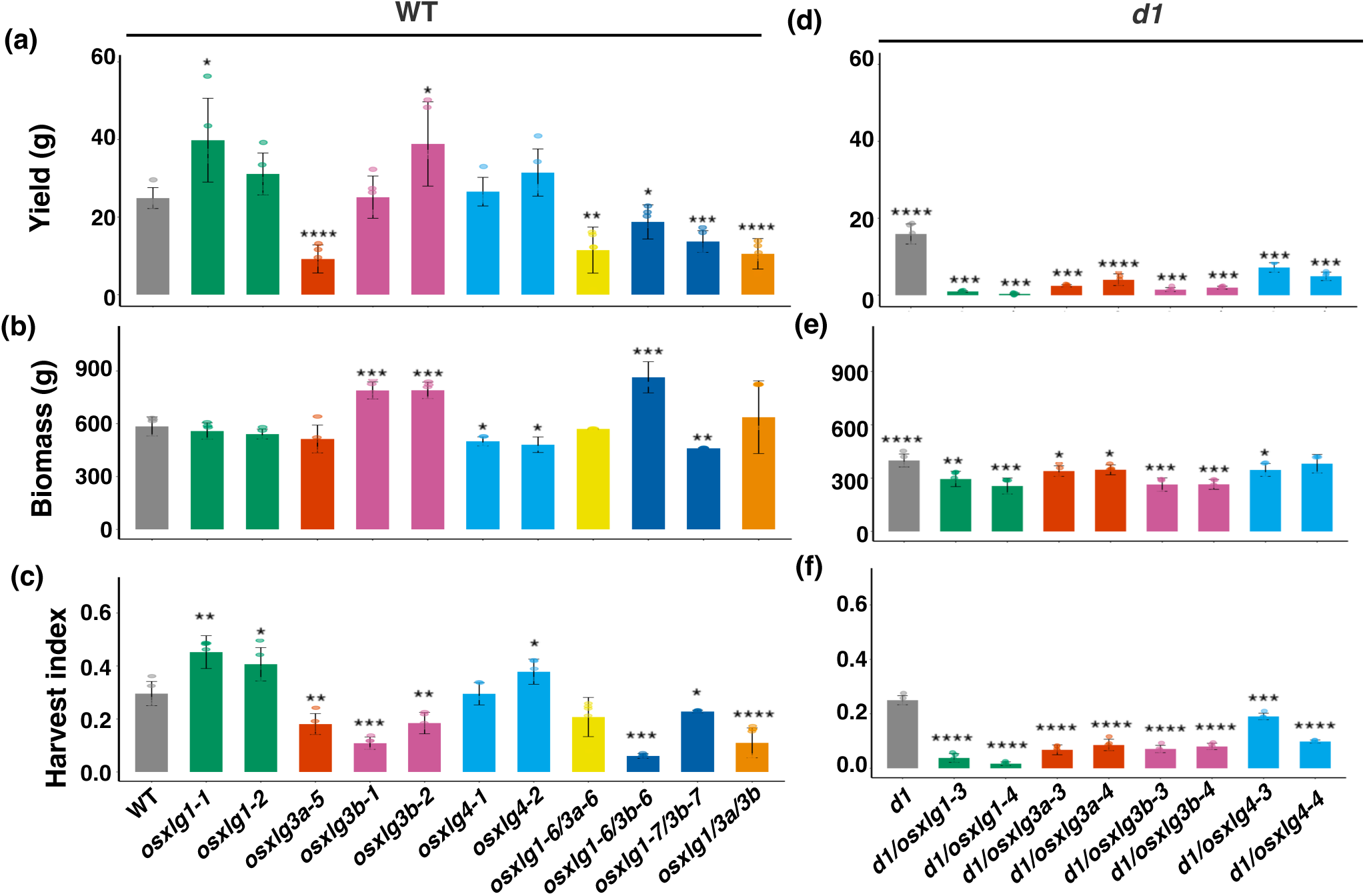
Effects of *OsXLG* mutations on plant performance. (a) Yield. (b). Above-ground biomass. (c) Harvest index. The *d1* mutant significant level was compared to WT. Statistical significance levels are indicated as follows: *, P < 0.05; **, P < 0.01; ***, P < 0.001, ****, P < 0.0001.

**Figure 7.**
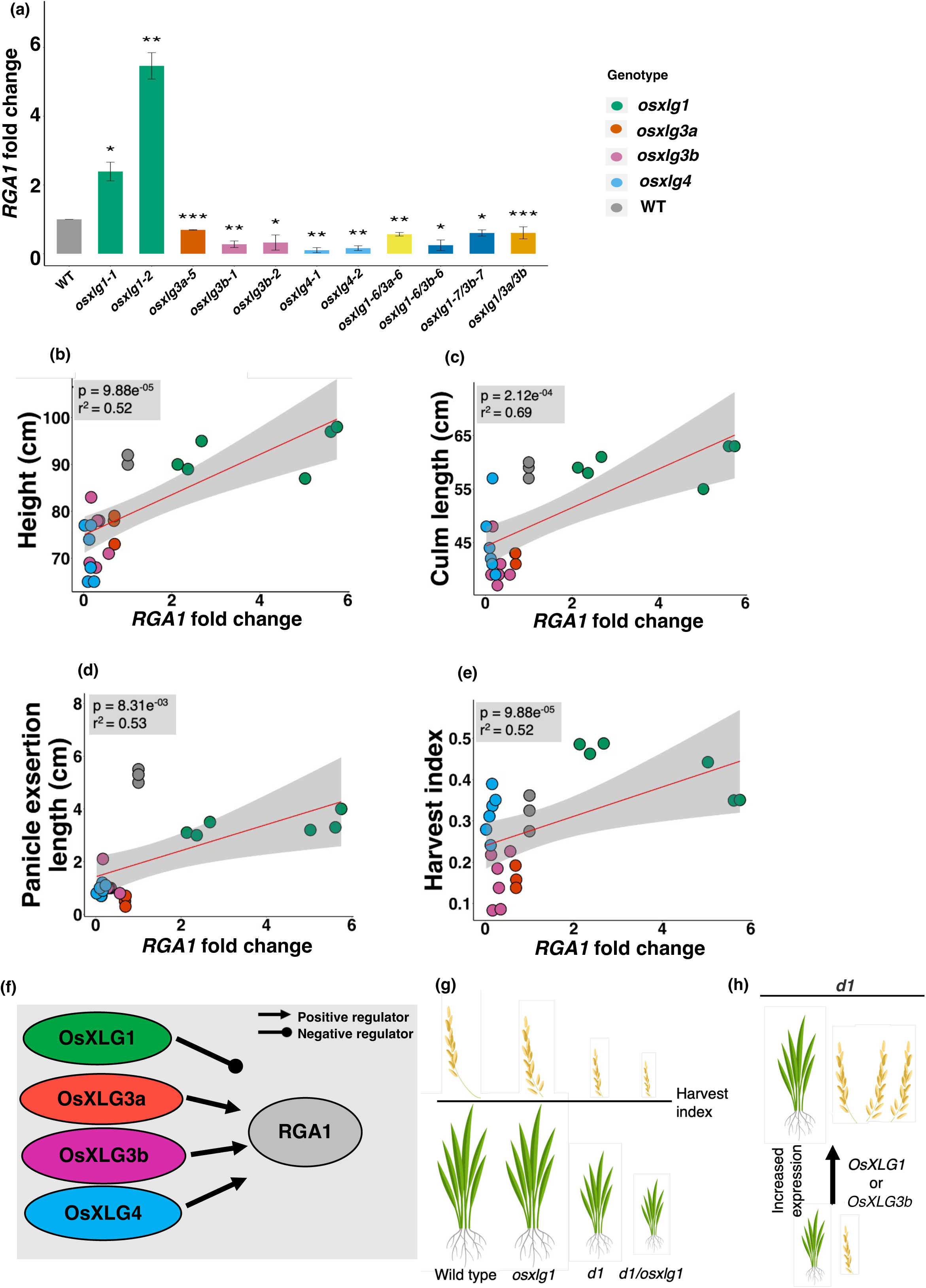
Analysis of *RGA1* expression in CRISPR mutants using qRT-PCR and correlation with various agronomic traits. Only agronomic traits showing significant correlations are depicted. (a) *RGA1* expression in *osxlg* CRISPR mutants. (b) Height. (c) Culm length (d) Panicle exsertion length. (e) Harvest index. (f) *RGA1* regulation by OsXLGs. (g) Effect of *OsXLG1* and/or *RGA1* mutations on plant architecture. (h) Proposed effect of *OsXLG1* and *OsXLG3a* expression on plant architecture. Statistical significance levels are indicated as follows: *, P < 0.05; **, P < 0.01; ***, P < 0.001, ****, P < 0.0001. The statistical relationships between *RGA1* expression and these traits were evaluated using linear regression, fitted with the ‘lm’ function in R. This analysis yielded p-values and correlation coefficients (R values), which quantitatively describe the significance and the degree of association between *RGA1* expression and each agronomic trait.

### CRISPR mutation impact on *OsXLG* expression

Figures 1c and 1d show results of qRT-PCR using primer pairs downstream of the CRISPR target site to determine the functional impact of CRISPR mutation on transcript abundance of the targeted *OsXLGs.* In both WT and *d1* mutant backgrounds, mutations in *OsXLG1* and *OsXLG3a* genes led to complete transcript absence (knockout), while mutations of *OsXLG3b* and *OsXLG4* genes resulted in reduced or undetectable transcript levels, depending on the allele.

Figures 2c to 2e illustrate the transcript levels of *osxlg* mutants received from Biswal et al. (2021). Mutations in the *OsXLG1* gene led to either reduced or undetectable transcript levels. Mutations in the *OsXLG3a* gene resulted in reduced or absent transcript levels, with the exception of *osxlg3a-7*. Mutations in the *OsXLG3b* gene consistently caused an absence of transcripts.

### Roles of OsXLGs with RGA1 in vegetative phenotypes

Figure 3 illustrates the effects of *OsXLG* mutations on vegetative traits. *osxlg1* mutants have a similar height as WT, while *osxlg3a, osxlg3b* and *osxlg4* mutants display shorter stature (Figure 3a, 3b). The reduced stature in *osxlg3a* and *osxlg3b* mutants primarily results from reduction in culm length, whereas *osxlg4* mutants exhibit reduction in both culm and flag leaf length (Figures 3c, 3d). Double and triple *osxlg* mutants also show shorter stature, primarily due to a reduction in culm length (Figures 3b, 3c). None of the *osxlg* mutants show differences in flag leaf width as compared to WT (Figures 3e). However, specific *osxlg* mutants display variation in tiller numbers: *osxlg3b* and *osxlg1-6/3b-6* mutants exhibit increased tiller numbers, whereas the *osxlg1-6/3a-6* mutant shows a decrease in tiller number (Figure 3f). *osxlg1*, *osxlg3a-5*, *osxlg4* and triple mutants maintain tiller numbers comparable to WT (Figure 3f). Interestingly, all *osxlg* mutants display early heading compared to WT (Figure 3g).

Previous studies (Ashikari *et al*., 1999; Oki *et al*., 2009; Peng *et al*., 2019; Yantong *et al*., 2022), along with our own results, show that *d1* mutant plants exhibit distinct vegetative characteristics when compared to the WT. These characteristics include shorter plant height (Figure 3b vs. 3i) due to reduced culm length (Figure 3c vs. 3j), and shorter and broader leaves (Figures 3d and 3e vs. 3k and 3l). Additionally, our study demonstrates that *d1* mutants maintain a similar number of tillers (Figure 3f vs. 3m) while exhibiting an earlier heading date compared to WT (Figure 3g vs. 3n). All *d1/osxlg* mutants exhibit even shorter stature than *d1* plants (Figures 3h, 3i). The decrease in height for *d1/osxlg4* mutants is due to reduced culm length (Figure 3j), for *d1/osxlg3a* and *d1/osxlg3b* mutants is due to reduced flag leaf length (Figure 3k), and for *d1/osxlg1* mutants arises from reductions in both (Figures 3j, 3k). The consistency in flag leaf width across all *d1/osxlg* mutants and the *d1* mutant (Figure 3l) indicates an independent role of *RGA1* in this trait (SI, Table 6). The reduction in tiller numbers in certain *osxlg* mutants in the *d1* but not the WT background, namely in *d1/osxlg1*, *d1/osxlg3a-3*, and *d1/osxlg4-3* (Figures 3f, 3m), implies redundancy of each of these *osxlgs* with *d1* for this trait. *d1* antagonizes *osxlg3b* in controlling tillering, as the presence of the *d1* mutation suppresses the increased tillering seen in the *osxlg3b* single mutant (Figure 3f). Moreover, the uniformity in heading dates among all *d1/osxlg* mutants and the *d1* mutant (Figure 3n), in contrast to the earlier heading observed for the *osxlg* mutants in the WT background (Figure 3g), supports an epistatic relationship with *d1*.

### Roles of OsXLGs with RGA1 in reproductive phenotypes

Figure 4 shows the effects of *OsXLG* mutations on reproductive traits. All single, double, and triple *osxlg* mutants exhibit shorter panicles compared to WT (Figure 4a, 4b). The panicle is composed of two sections: the length from the panicle base to the first spikelet (the panicle exsertion length (PEL)), and the length of the main stem of the panicle from the first spikelet to the tip (panicle axis length (PAL)) (see labels in Figure 4a). Interestingly, the shorter panicles of all *osxlg* mutants result from reduction in PEL (Figure 4c), while *osxlg3a-5*, *osxlg3b-1*, *osxlg4*, and double and triple *osxlg* mutants additionally show a decreased PAL (Figures 4d). In contrast, *osxlg1* and *osxlg3b-2* mutants maintained a PAL comparable to WT (Figure 4d). Notably, *osxlg3b,* and *osxlg1-6/3b-6* mutants produce more panicles while *osxlg1-6/3a-6* mutant produce fewer panicles (Figure 4e); the other mutants display similar panicle number as WT (Figure 4e). Spikelets per panicle, grains per panicle and the spikelet fertility — which is the proportion of total number of grains per panicle/total number of spikelet per panicle — were also measured. *osxlg1* and *osxlg4* mutants show an increase in both spikelets per panicle and grains per panicle compared to WT (Figure 4f, 4g). On the other hand, *osxlg3a-5*, *osxlg3b-1*, *osxlg1/3b*, and the triple mutants exhibit a reduction in these parameters compared to WT (Figure 4f, 4g). The *osxlg1-6/3a-6* mutant produces a comparable number of spikelets per panicle but with fewer grains per panicle as compared to WT (Figure 4f, 4g). Additionally, spikelet fertility is highly reduced in *osxlg3a-5*, *osxlg3b*, *osxlg1/3b* and triple mutants, while *osxlg1, osxlg4* and *osxlg1-6/3a-6* exhibit similar spikelet fertility as WT (Figure 4h).

Expanding on previous studies that *d1* mutants possess shorter panicles (Ashikari *et al*., 1999; Oki *et al*., 2009), our study further explored this feature by measuring the PEL and PAL (see labels in Figure 4i). The observed shorter panicles (Figure 4b vs. 4j) in *d1* mutants can be attributed to decreases in both PEL and PAL (Figures 4c and 4d vs. 4k and 4l). Our results also show that *d1* mutants exhibited an increase in the number of spikelets (Figure 4f vs. 4n) and grains per panicle (Figure 4g vs. 4o), with panicle numbers (Figure 4e vs. 4m) and spikelet fertility similar to WT (Figure 4h vs. 4p). However, remarkably, combining *OsXLG* mutations with the *d1* mutation led to a detrimental impact on all of these traits (Figure 4j, n, o). This is despite the fact that in the WT background, only *osxlg3a* and *osxlg3b* mutants showed reductions in these traits. The further decreased counts for these traits for *d1/osxlg3a* and *d1/osxlg3b* mutants suggest that *d1* has an enhancing effect on *osxlg3a* or *osxlg3b.* In contrast, the reduced counts for these traits for *d1/osxlg1* and *d1/osxlg4* mutants exhibit a mutual antagonistic effect between *osxlg1* or *osxlg4* with *d1*, where their individual mutations mutants display increased counts for these traits (Figures 4f, 4g). In terms of spikelet fertility, all *d1/osxlg* mutants except the *d1/osxlg4-3* mutant showed decreased fertility relative to *d1* (Figure p).

### Roles of OsXLGs with RGA1 in grain and post-harvest phenotypes

Figure 5 illustrates the grain length and width, and the 100-grain weight of the different genotypes. In the WT background, *osxlg3b-1, osxlg4, osxlg1-6/3b-6* mutants display reduced grain length, while the *osxlg1-6/3a-6* mutant shows an increase in grain length (Figure 5a, 5b). *osxlg1* and *osxlg3a-5* mutants have similar grain length compared to WT (Figure 5a, 5b). Notably, all *osxlg* mutants exhibit similar grain width to WT (Figure 5a, 5c). *osxlg3b-1, osxlg4*, *osxlg1/osxlg3b* mutants as well as triple mutants show lower 100-grain weight, while *osxlg1*, *osxlg3a*, and *osxlg1-6/3a-6* mutants show similar 100-grain weight to WT (Figure 5c).

*d1* mutants exhibit shorter (Figure 5a and 5b vs. 5e and 5f) but not narrower (Figure 5c vs. 5g) seeds than WT, leading to a round seed phenotype, which in turn leads to reduced grain weight (Figure 5d vs. 5h). The *d1/osxlg* mutants show further reduction in grain length, except the *d1/osxlg4-4* mutant, which has similar grain length as the *d1* mutant (Figure 5f). This suggests that *osxlg1*, *osxlg3a* and *osxlg3b* have an enhancing effect on *d1*, while *osxlg4* and *d1* are partially redundant. While none of the *osxlg* mutations alter grain width in the WT background, *d1/osxlg3a* and *d1/osxlg3b-3* mutants produce narrower grains than *d1* (Figure 5g), suggesting that *osxlg3a* and *osxlg3b* each exhibit redundancy with *d1* for grain width. Furthermore, *d1* mutants have reduced 100-grain weight and all *d1/osxlg* mutants exhibit a further reduction in grain weight (Figure 5h).

Figure 6 displays the total yield, dry shoot biomass, and harvest index of the *osxlg* mutants. In the WT background, *osxlg3a-5*, *osxlg1-6/3a-6*, *osxlg1/3b*, and the triple mutants exhibit a lower yield than WT (Figure 6a), whereas *osxlg1-1* and *osxlg3b-2* single mutants show a higher yield compared to WT (Figure 6a). *osxlg4* mutants have similar yield to WT (Figure 6a). In terms of dry shoot biomass, *osxlg3b* and *osxlg1-6/3b-6* mutants have higher biomass (Figure 6b), while *osxlg4* and *osxlg1-7/3b-7* mutants have lower biomass (Figure 6b), while *osxlg1* and *osxlg1-6/3a-6* mutants display shoot biomass similar to WT (Figure 6b). Harvest index (HI) is calculated as the grain yield (g) to total dry shoot biomass (g) ratio. In this context, both *osxlg1* mutants as well as *osxlg4-2* mutants outperform WT (Figure 6c). For *osxlg1* mutants, the higher HI is attributed to increased yield with similar shoot biomass as WT, whereas for the *osxlg4-2* mutant the higher HI is attributable to lower shoot biomass while maintaining yield comparable to WT. In contrast, *osxlg3a, osxlg3b, osxlg1/3b*, and triple mutants demonstrate reduced HI (Figure 6c), mainly because of significant reductions in total yield. However, the *osxlg1-6/3a-6* mutant shows a HI similar to WT (Figure 6c).

The *d1* mutants in our study exhibit both a lower yield (Figure 6a vs. 6d) and reduced biomass (Figure 6b vs. 6e) than WT; this results in a harvest index (HI) comparable to WT (Figure 6c vs. 6f). In contrast to favorable impacts on yield and HI of some *osxlg* mutations in the WT background, *osxlg* mutations in the *d1* background have a uniformly deleterious impact on these traits (SI, Table 7). That *osxlg* mutations exacerbate the already negative impacts of *d1* mutation on yield and harvest index (Figure 6d, 6e) is particularly evident for the *d1/osxlg1* mutants, where average yield was 0.64 grams, as compared to 15.9 grams for *d1*. As a result, the *d1/osxlg* double mutants had a significantly lower HI (Figure 6f) than *d1*, despite the fact that these double mutants (except for the *d1/osxlg4-4* mutant), also demonstrate a reduced dry shoot biomass as compared to *d1* (Figure 6e).

### Dynamic Interplay Between RGA1 and OsXLGs in Gene Expression and Phenotypic Correlation

To unravel the complexity of Gα interplay, we quantified the relative transcript levels of non-CRISPR targeted *OsXLGs* and *RGA1* in each *osxlg* mutant in the WT background.

We observed a consistent reduction in *RGA1* expression in all CRISPR mutants, with the exception of *osxlg1* mutants, which displayed *RGA1* upregulation (Figure 7a). We then conducted correlation analyses to investigate potential relationships between *osxlg* single mutant phenotypes and *RGA1* expression. SI, Table 5 provides r^2^ and p values of these correlations for all phenotypic traits. Our results reveal a positive correlation between *RGA1* expression and two key vegetative traits: plant height and culm length (Figures 7b, 7c). Our results also reveal a positive correlation between *RGA1* expression and key reproductive traits: panicle exsertion length and harvest index (HI) (Figures 7d, 7e).

Conversely, to explore the impact of the absence of RGA1 on *OsXLG* gene expression, we quantified the transcript levels of non-CRISPR targeted *OsXLG* genes in the WT and *d1/osxlg* mutants. *OsXLG* genes exhibited complex differential regulation across various genetic backgrounds, with patterns of upregulation and downregulation varying depending on the specific *osxlg* mutant combinations (SI, Figure 4).

We also conducted an analysis to investigate the relationship between the expression of *OsXLG* genes and all of the measured agronomic traits in both the WT and *d1* mutant backgrounds. The results of these correlations are presented in SI, Table 5, including the r^2^ and p-values for each phenotype. Notably, a greater number of significant correlations between *OsXLG* transcript abundance and phenotypes are seen in the *d1* than in the WT background (SI, Table 6). In particular, in the *d1* mutant background, *OsXLG1* expression is positively correlated with the key agronomic traits plant height and culm length (SI, Figure 5a, 5b), panicle and axis length (SI, Figure 5c, 5d), spikelet and grain counts per panicle (SI, Figure 5e and 5f), fertility, and harvest index (SI, Figure 5g, 5h), while *OsXLG3b* expression is positively associated with tiller number (SI, Figure 5i), flag leaf dimensions (SI, Figure 5j, 5k), panicle number and grains per panicle (SI, Figure 5l, 5m), fertility and biomass (SI, Figure 5n 5o), yield, and harvest index (SI, Figure 5p and 5q).

## Discussion

Leveraging CRISPR gene-editing technology, we generated new alleles of *OsXLGs* in both Nipponbare wild type and in a *d1* mutant in the Nipponbare background, aiming to decipher the interplay between the canonical Gα, RGA1, and the non-canonical OsXLGs in control of plant architecture and productivity. This approach yielded a range of genetic variants (Figures 1a, 1b). Unlike previous studies (Biswal *et al*., 2021; Cui *et al*., 2020; Zhao *et al*., 2022), we identified CRISPR-free, homozygous T2 lines, and we quantified the impact of each edit on transcript abundance (Figures 1c, 1d). For *OsXLG1* and *OsXLG3a*, the lines were essentially transcript null mutants, for *OsXLG3b* the extent of suppression differed across various lines, and for *OsXLG4* all edits resulted in transcript knockdown. We also went beyond the scope of previous studies (Biswal *et al*., 2021; Cui *et al*., 2020; Zhao *et al*., 2022) by examining the role of OsXLGs across the entire spectrum of plant architecture, spanning vegetative, reproductive, and post-harvest stages for a total of 19 traits.

We found that single Gα mutants had marked impacts on agronomic traits. Several consistent patterns emerge from the phenotypes of *osxlg* mutants. Notably, traits such as heading date (Figure 3g), panicle length (Figure 4b), and panicle exsertion length (Figure 4c) consistently decrease in all *osxlg* mutants in the WT background, effects also observed in the *d1* null mutant of the canonical Gα, *RGA1* (Figures 3n, 4j, 4k, and Table 1). Heading date, in particular, is a crucial agronomic trait because it influences the timing of flowering, adaptation to environmental conditions, and overall yield (Lin *et al*., 2021). Moreover, Arabidopsis XLG2 proteins influence flowering by interacting with transcriptional regulators like RELATED TO VERNALIZATION1 (RTV1). RTV1, a DNA-binding protein, regulates key flowering genes such as SUPPRESSOR OF OVEREXPRESSION CONSTANS1 (SOC1) and FLOWERING LOCUS T (FT) (Heo *et al*., 2012). XLG2 enhances RTV1’s DNA-binding activity, particularly when bound to GTP, leading to the upregulation of these floral integrator genes and early flowering (Heo *et al*., 2012). Gα proteins in rice may similarly interact with transcriptional regulators to modulate gene expression during critical developmental processes, such as the transition to flowering.

Flag leaf width is broadened in the *d1* mutant but remains unaltered in *osxlg* mutants (Figure 3e) (Figure 3l, Table 1). This alteration, observed solely in the *d1* mutant, could be attributed to a disruption of RGA1-Gβγ signaling that is not compensated for by the OsXLG proteins. We also observed gene-specific effects of OsXLG proteins. In the WT background, *osxlg1* mutant plants, despite being transcript nulls (Figure 1c, Figure 7g), were more phenotypically WT than the other *osxlg* mutants. For example, in *osxlg1* mutants alone, traits such as height, culm length, and panicle axis length did not display significant deviations from the wild type (Figures 3b, 3c, 4d, and Table 1). These results suggest that OsXLG1 may have a unique and specific role, distinct from the other OsXLGs or RGA1.

Our gene expression analyses further implicate a unique role of OsXLG1. Our findings show that, among the OsXLGs, only OsXLG1 functions as a negative regulator of *RGA1* transcript abundance: mutations in *OsXLG1* lead to plants with increased expression of *RGA1* (Figure 7a, 7f). Increased *RGA1* expression in *osxlg1* null mutants in the WT background (Figure 7a) is associated with favorable reproductive traits, including longer grains, more yield, higher grain weight and harvest index (Figure 7g, Table 2) and unfavorable vegetative traits such as increased height, and culm length, and higher biomass (Figure 7g, Table 2). All of these phenotypes are oppositely impacted in the *d1* vs the *d1/osxlg1* mutants. Further investigation is needed to determine the mechanism underlying OsXLG1 regulation of *RGA1* expression.

**Table 2.** Summary of observed effects of Gα mutations on rice agronomic traits. Designations are relative to WT for *d1* and relative to the genotypic background (WT or *d1*) for the *OsXLG* edited lines, as indicated. A1- Allele 1, A2- Allele 2, A3- Alle 3, A4- Allele 4 designations as in Figure 1b.

| Phenotype | RGA1 | OsXLG1 |  | OsXLG3a |  | OsXLG3b |  | OsXLG4 |  |
| --- | --- | --- | --- | --- | --- | --- | --- | --- | --- |
| Background<br>→ | *d1* | WT | *d1* | WT | *d1* | WT | *d1* | WT | *d1* |
| Plant height | Decreased | No change | Decreased | Decreased | Decreased | Decreased | Decreased | Decreased | Decreased |
| Culm length | Decreased | No change | Decreased | Decreased | No change | Decreased | No change | Decreased | Decreased |
| Flag leaf length | Decreased | No change | Decreased | No change | A3-Decreased† | Decreased | Decreased | Decreased | No change |
| Flag leaf width | Decreased | No change | No change | No change | No change | No change | No change | No change | No change |
| Tiller number | No change | No change | *Decreased* | No change | *Decreased* | Increased | *Decreased* | No change | A4-<br>*Decreased*† |
| Days to heading | Decreased | Decreased | No change | Decreased | No change | Decreased | No change | Decreased | No change |
| Panicle length | *Decreased* | *Decreased* | *Decreased* | *Decreased* | *Decreased* | *Decreased* | *Decreased* | *Decreased* | *Decreased* |
| Panicle exertion length | *Decreased* | *Decreased* | No change | *Decreased* | No change | *Decreased* | No change | *Decreased* | No change |
| Panicle axis length | *Decreased* | No change | *Decreased* | *Decreased* | *Decreased* | A1-<br>*Decreased*† | *Decreased* | *Decreased* | *Decreased* |
| Panicle number | No change | No change | *Decreased* | No change | *Decreased* | Increased | *Decreased* | No change | A4-<br>*Decreased*† |
| Spikelets per panicle | Increased | Increased | *Decreased* | *Decreased* | *Decreased* | *Decreased* | *Decreased* | Increased | *Decreased* |
| Grains per panicle | Increased | Increased | *Decreased* | *Decreased* | *Decreased* | *Decreased* | *Decreased* | Increased | *Decreased* |
| Spikelet fertility | No change | No change | *Decreased* | *Decreased* | *Decreased* | *Decreased* | *Decreased* | No change | *Decreased* |
| Grain length | *Decreased* | No change | *Decreased* | No change | *Decreased* | *A1-*<br>*Decreased†* | *Decreased* | *Decreased* | *A3-*<br>*Decreased†* |
| Grain width | No change | No change | No change | No change | *Decreased* | No change | *A4-*<br>*Decreased†* | No change | No change |
| 100- grain weight | *Decreased* | No change | *Decreased* | No change | *Decreased* | *A2-*<br>*Decreased†* | *Decreased* | *Decreased* | *Decreased* |
| Yield | *Decreased* | *A1-*<br>*Increased†* | *Decreased* | *Decreased* | *Decreased* | *A1-*<br>*Increased†* | *Decreased* | No change | *Decreased* |
| Biomass | *Decreased* | No change | *Decreased* | No change | *Decreased* | *Increased* | *Decreased* | *Decreased* | *A3-*<br>*Decreased†* |
| Harvest index | No change | Increased | *Decreased* | *Decreased* | *Decreased* | *Decreased* | *Decreased* | Increased* | *Decreased* |

†- Only one allele showed effect

By contrast, height, culm length, and panicle axis length were all decreased across *osxlg3a, osxlg3b*, *osxlg4* and *d1* (Figures 3i, 3j, 4l, and Table 1), suggesting that for these traits OsXLG3a, OsXLG3b, and OsXLG4 might function in pathways that closely parallel the RGA1-Gβγ signaling pathway. Moreover, OsXLG3a, OsXLG3b, and OsXLG4 appear to play a role in positively regulating *RGA1* expression, as mutations in these genes result in plants with decreased *RGA1* expression (Figures 7a, 7f). The similar phenotypes observed in *osxlg3a*, *osxlg3b*, *osxlg4*, and *d1* mutants (Table 2), along with their shared characteristic of reduced *RGA1* expression, suggest two possible hypotheses. First, there may be a shared regulation at the transcriptional level, where these genes collectively influence *RGA1* expression. Yantong et al. (2022) identified rice cultivar carrying a weak allele of *RGA1*, leading to reduced *RGA1* expression levels (Yantong *et al*., 2022). These rice plants exhibit shorter stature, dense and erect panicles, smaller seeds, but with more grains per panicle. Consistent with their observations and considering *RGA1* expression levels across all single mutant backgrounds, we observe that *RGA1* abundance is significantly positively correlated with height (Figure 7b), culm length (Figure 7c), panicle exsertion (Figure 7d), and harvest index (Figures 7e). Second, their similar impact on these traits implies that, in *d1*, these OsXLGs may compensate for the loss of RGA1 function at the protein level, either by signaling to RGA1’s downstream partners or by substituting for RGA1 in regulation of Gα -Gβγ signaling. Indeed, in our recent study (Ferrero-Serrano *et al*., 2024), we found that physical interactions with the five Gβγ dimers of rice are similar for RGA1, OsXLG3a, OsXLG3b, and OsXLG4, while OsXLG1 interacts only with particular Gβγ combinations; Arabidopsis XLGs also exhibit differential specificity in Gβγ interaction (Ferrero-Serrano *et al*., 2024). This reinforces the notion that OsXLG1 operates in a distinct and separate manner from RGA1 and the other OsXLGs.

Our own *osxlg3a* mutants did not survive beyond the T1 generation and exhibited compromised growth (SI, Figure 2a). We sequenced the potential CRISPR off-target region with the highest sequence similarity to the *OsXLG3a’s* target site and observed no mutations (SI, Figure 2b), indicating that the lethality is likely due to the *OsXLG3a* mutation. However, a CRISPR *osxlg3a-5* mutant from Biswal et al. (2021) did not display compromised growth in our conditions and exhibited phenotypes consistent with their previous report (Biswal et al., 2021). Our *osxlg3a* mutant is predicted to truncate OsXLG3a (Figure 1b), in contrast to Biswal’s *osxlg3a* mutant, which is predicted to eliminate essentially the entire protein (Figure 2b), suggesting a possible dominant negative effect of the OsXLG3a N-terminus in our mutant. By contrast, our mutation of *OsXLG3a* in the *d1* background was not lethal, (Figure 3h), despite the predicted truncation being similar to that in the WT background. Compensatory mechanisms triggered by the *d1* mutation might alleviate the deleterious effects of the *OsXLG3a* mutation. These contrasting phenotypes emphasize that further investigations are warranted to elucidate the as yet unknown roles of the N-terminal domain in OsXLG function. In Arabidopsis, XLG2 and XLG3 contain nuclear localization signals within the N-terminal domain indicating a potential role in nuclear localization (Chakravorty *et al*., 2015).

Like *osxlg3a*, *osxlg3b* mutants also exhibit decreased spikelets (Figure 4f) and grains per panicle (Figure 4g), and lower fertility (Figure 4h) resulting in a lower harvest index (Figure 6c). This similarity in their phenotypic profile (Table 1) suggests that OsXLG3a and OsXLG3b likely share overlapping functions. This shared pattern may be a result of the recent gene duplication event in monocots (Cantos *et al*., 2023; Mohanasundaram *et al*., 2022) that led to the formation of *OsXLG3a* and *OsXLG3b*, allowing them to retain similar functions.

Our recent phylogenetic analysis identified OsXLG4 as a legitimate XLG family member. Our current study reveals distinct traits of *osxlg4* mutants (Table 2), confirming that previous designation of this gene as a non-functional pseudogene (Mohanasundaram *et al*., 2022) was incorrect. There is an intriguing contrast in the phenotypic outcomes of *osxlg4* mutants in WT and *d1* backgrounds (Table 2). In the WT background, *osxlg4* mutants exhibit an increase in spikelets and grains per panicle (Figures 4g, 4h), while in the *d1* background, they display the opposite (Figures 4o, 4p, Table 2). These antagonistic effects suggest that the presence of RGA1 influences the role of OsXLG4 in these aspects. Interestingly, the expression of *OsXLG3a* and *OsXLG3b* shows opposite patterns in *osxlg4* vs. *d1/osxlg4* mutants, suggesting that these genes might contribute to the opposing phenotypic effects of *OsXLG4* mutation in the WT vs. *d1* backgrounds.

Our findings on *osxlg* single mutant phenotypes both align with and diverge from previous studies (Biswal *et al*., 2021; Cui *et al*., 2020; Zhao *et al*., 2022). Results that diverge from previous reports could be due to several factors, including possible residual CRISPR components in other studies, variation in the functionality of differentially truncated XLG proteins, and diverse growth conditions. Biswal et al.’s study (2021) showed contrasting phenotypes in the same mutants when grown under the different environmental conditions of tropical versus temperate greenhouses (Biswal *et al*., 2021). We also observed an impact of growth conditions on mutant phenotypes, supporting genotype by environment interactions. For example, the *osxlg3a-5* mutant generated by Biswal et al. (2021) in our hands exhibited no changes in the number of tillers, grain length and biomass (Table 2), opposite of the increases reported by Biswal et al. (2021). We also observed no change in panicle number and grain weight, opposite of their reported decreases. Such findings emphasize the significance of environmental factors in influencing the impact of genetic modifications on rice agronomic traits. However, our phenotypic results for the *osxlg* double and triple mutants generated by Biswal et al. (2021) generally paralleled their findings. It is also worth noting that Cui et al. (2020), which was the only previous study to generate *OsXLG4* CRISPR alleles (referred to as *pxlg3*), reported different phenotypes than ours for height, grains per panicle and 100-grain weight (Table 2). While our *osxlg4* mutants are transcript knockdowns, the impact of Cui’s *OsXLG4* mutations on gene expression was not reported.

We additionally found that double, and triple *OsXLG* mutations, as well as *OsXLG* mutant combinations with the *d1* mutation, significantly affect rice architecture and, importantly, the overall yield and harvest index (Table 2). By comparing double, and triple mutants with single mutants, we uncovered patterns of partial redundancy and epistasis among the OsXLGs (SI, Table 7). Our analysis suggests that OsXLGs in some instances compensate for each other’s loss and the redundancy adds another layer of genetic security for some key agronomic traits, e.g., height, yield, and harvest index (double and triple *osxlg* mutants; Figures 3a, 6a, 6c). In Arabidopsis as well, T-DNA insertional mutants of *XLGs* display functional redundancy in some traits, e.g., in root morphogenesis (Ding *et al*., 2008) and in PAMP-triggered MAPK signaling and immune response (Wang *et al*., 2023). Intricate relationships between OsXLGs and RGA1 are also evident in their genetic interactions in controlling overall plant architecture (Table 2; SI, Table 6). Similar results were also observed in the study on maize Gα proteins (Wu *et al*., 2018) revealing partial redundancy between canonical Gα, CT2, and maize XLGs in controlling SAM size and height, but an independent role of CT2 in ear fasciation. Moreover, in the absence of RGA1 in the *d1* mutant background, the positive correlations observed between *OsXLG1* and *OsXLG3b* abundance and multiple agronomic traits (Figure 7h; Si Table 5; SI Figure 5) suggest that these non-canonical Gα proteins play a crucial role in maintaining growth and yield, ensuring the plant continues to develop effectively despite the disruption of canonical Gα signaling (Figure 7h).

In summary, our study reveals an intricate interplay between OsXLGs and RGA1 in the regulation of Gα subunit expression, rice architecture and yield. The roles of OsXLG proteins in both intra-family interactions and their interactions with RGA1 hold significant implications for crop improvement. Harnessing the inherent redundancy and synergistic potential within these genetic networks could pave the way for developing rice varieties with enhanced growth and yield, ultimately advancing agricultural resilience and productivity. For example, upregulation of gene expression through promoter editing of the *RGA1, OsXLG3a, OsXLG3b* genes may enhance yield. Moreover, the overexpression of *OsXLG3a* or *OsXLG3b* specifically in the panicle could contribute to an increase in panicle length and exsertion, spikelets per panicle, and grains per panicle. This integrated perspective on OsXLG and RGA1 interactions opens up new avenues for precision breeding and crop improvement in rice, ultimately aiming to address global food security challenges.

## Acknowledgements

This research was supported by USDA/NIFA grant number: 2019-67013-29234 to SMA. CFC gratefully acknowledges support from the Penn State Intercollege Graduate Degree Program in Plant Biology, and from a Penn State Henry W. Popp Fellowship. We thank Drs. Alan M. Jones and Akshaya Biswal for generously sharing their *XLG* edited rice lines. We also thank Dr. Jiang Kang Zhu for providing the pSSTU CRISPR constructs.

