## Supplemental Information for "Manipulating rice canonical Gα and extra-large G protein subunits for improved agronomic traits"

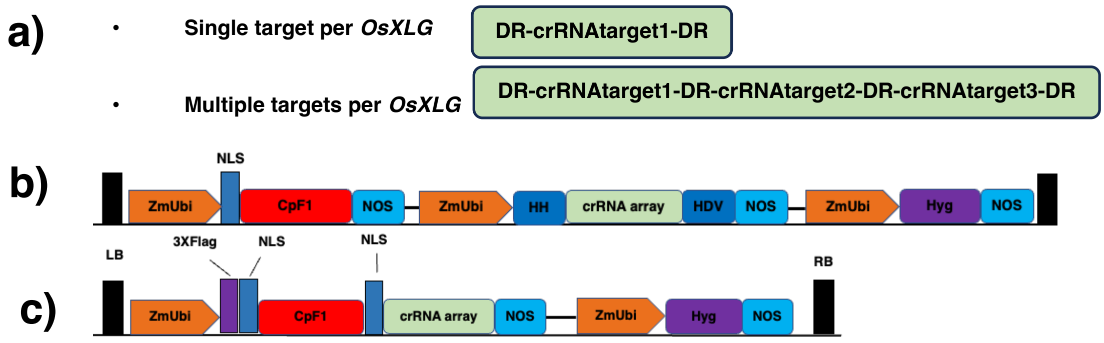


**SI, Figure 1. Overview of CRISPR-Cpf1 targeting strategies and systems in *OsXLG* genes manipulation.** a) Strategies employed in *OsXLG* editing, encompassing two specific approaches: (1) a single target strategy for each *OsXLG* gene (2) a multiple target strategy for each *OsXLG* gene. The figure also includes detailed schematic diagrams of two CRISPR systems: (b) the pCRISPR_ribozyme system, highlighting components such as the *ZmUBI1* promoter, LbCpf1 nuclease, and various ribozyme elements, adapted from Tang et al., 2018; and (c) the pCRISPR_SSTU system, illustrating key elements like the *ZmUBI* promoter with a 3xFlag tag, Cpf1 nuclease, and the CRISPR RNA target site array, modified from Wang et al., 2018.

**(a)** **CRISPR-*OsXLG3a* transformation**

| Gene | No. of plants generated | No. of plants with mutation | Mutation type |
|---|---|---|---|
| *OsXLG3a* | 5 | 5 | Deletions |

**(b)** **Mutation identification**

```
Sequence alignment*
WT      TTTCTTCCCTGCCGCAACGCGAGAATACGCTC
Allele1 TTT-------------------------GCTC (-25bp)
Allele2 TTTCTTCCCTGCCGCAACGC----------TC (-10bp)
```

*All generated plants have the same mutation type

**(c)** **Developmental stages**

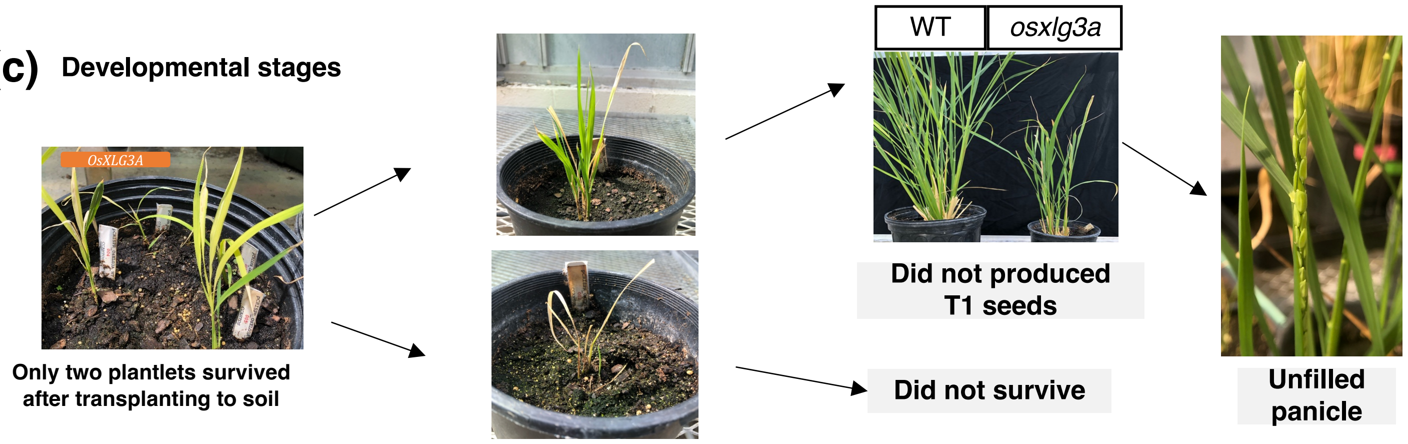


**(d)** **Off target analysis**

**LOC_Os02g57820 genomic, AT hook motif domain containing protein, expressed**

Expect = 0.10 Identities = 17/17 (100%)

```
crRNA_osxlg3a: 1             tttccatagttcctcaa 17
                             |||||||||||||||||
potential_offtarget: 2966    tttccatagttcctcaa 2982
```

**(e)** **Sequence alignment of potential off-target site**

```
LOC_Os02g57820  TTTTCCATAGTTCCTCAATTGAGTAAAAGGCTTTGATGCTGCTTTTAGATGAACTTTTGC  360
PCR_sequence    TTTTCCATAGTTCCTCAATTGAGTAAAAGGCTTTGATGCTGCTTTTAGATGAACTTTTGC  360
                ************************************************************
```

**SI, Figure 2. Compromised growth of *osxlg3a* mutants.** (a) Rice transformation summary of CRISPR-*osxlg3a* in the WT background. (b) Mutation types identified in *osxlg3a* mutants. (c) Effects of *osxlg3a* mutation on growth and development. (d) BLAST analysis to identify potential off-target sites of CRISPR. (e) Sequence analysis of the potential off-target site.

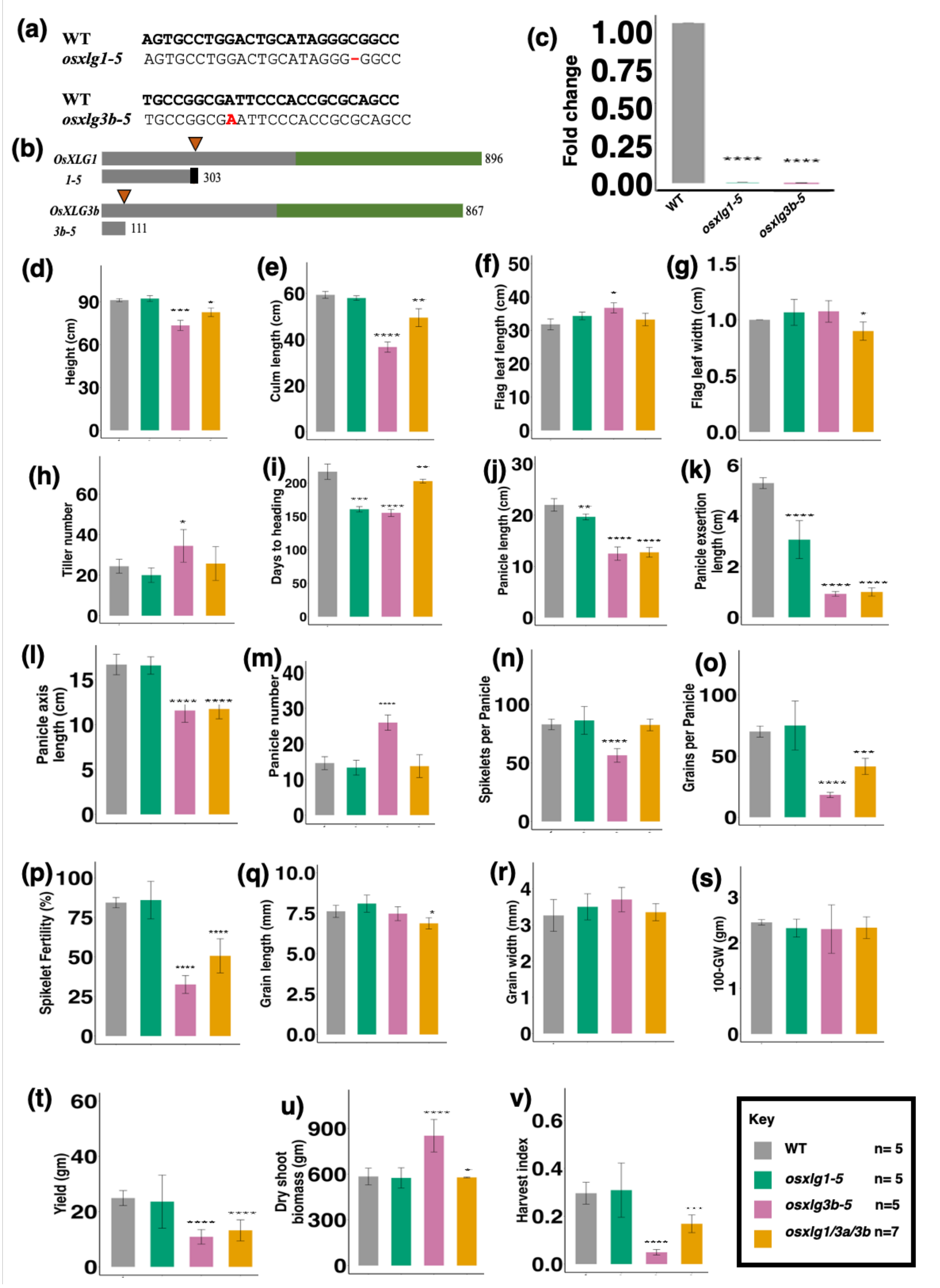


**SI, Figure 3. Non-CRISPR free mutants from Biswal et al. (2021).** All of these CRISPR *osxlg* alleles are in the WT background. The triple *osxlg* mutant phenotypes are consolidated data from all non CRISPR-free lines of the three genotypes of *osxlg1/3a/3b*. (a) Sequence alignments of *osxlg1-5* and *osxlg3b-5*. (b) Schematic representation of the OsXLG proteins and the predicted proteins of *osxlg1-5* and *osxlg3b-5*. The orange arrowheads indicate the position of the CRISPR RNA target sites the numbers indicate the lengths of the corresponding amino acid sequences. Grey bars-N-terminal domain; Green bars-Gα domain; Black bars- position of the frameshift mutations and amino acid deletions. (c) Relative expression levels (fold change) of *osxlg1-5* and *osxlg3b-5*determined by quantitative real-time polymerase chain reaction (qRT-PCR), as compared to control (unedited) plants. (d-v). Mutant phenotypes of non-CRISPR free *osxlg* mutants. Statistical significance is indicated as follows: *, $P<0.05$; **, $P<0.01$; ***, $P<0.001$; ****, $P<0.0001$. The p-values were generated using Student's t-tests to assess the statistical significance of the observed differences between the mutant and control groups. For non-CRISPR-free *osxlg1-5* mutants, spikelet and grains per panicle exhibit a decrease, contrary to CRISPR-free *osxlg1* mutants, resulting in lower yield and harvest index, which is the opposite of CRISPR-free *osxlg1* mutants. Similarly, non-CRISPR-free *osxlg3b-5* mutants show inconsistent phenotypic outcomes, including flag leaf length, grain weight and length, and yield, compared to CRISPR-free *osxlg3b* mutants. Finally, the consolidated non-CRISPR-free *osxlg1/osxlg3a/osxlg3b* mutants exhibit inconsistent phenotypic results, such as flag leaf width, spikelets per panicle, grain weight and length, and biomass, compared to the consolidated CRISPR-free *osxlg1/osxlg3a/osxlg3b* triple mutants.

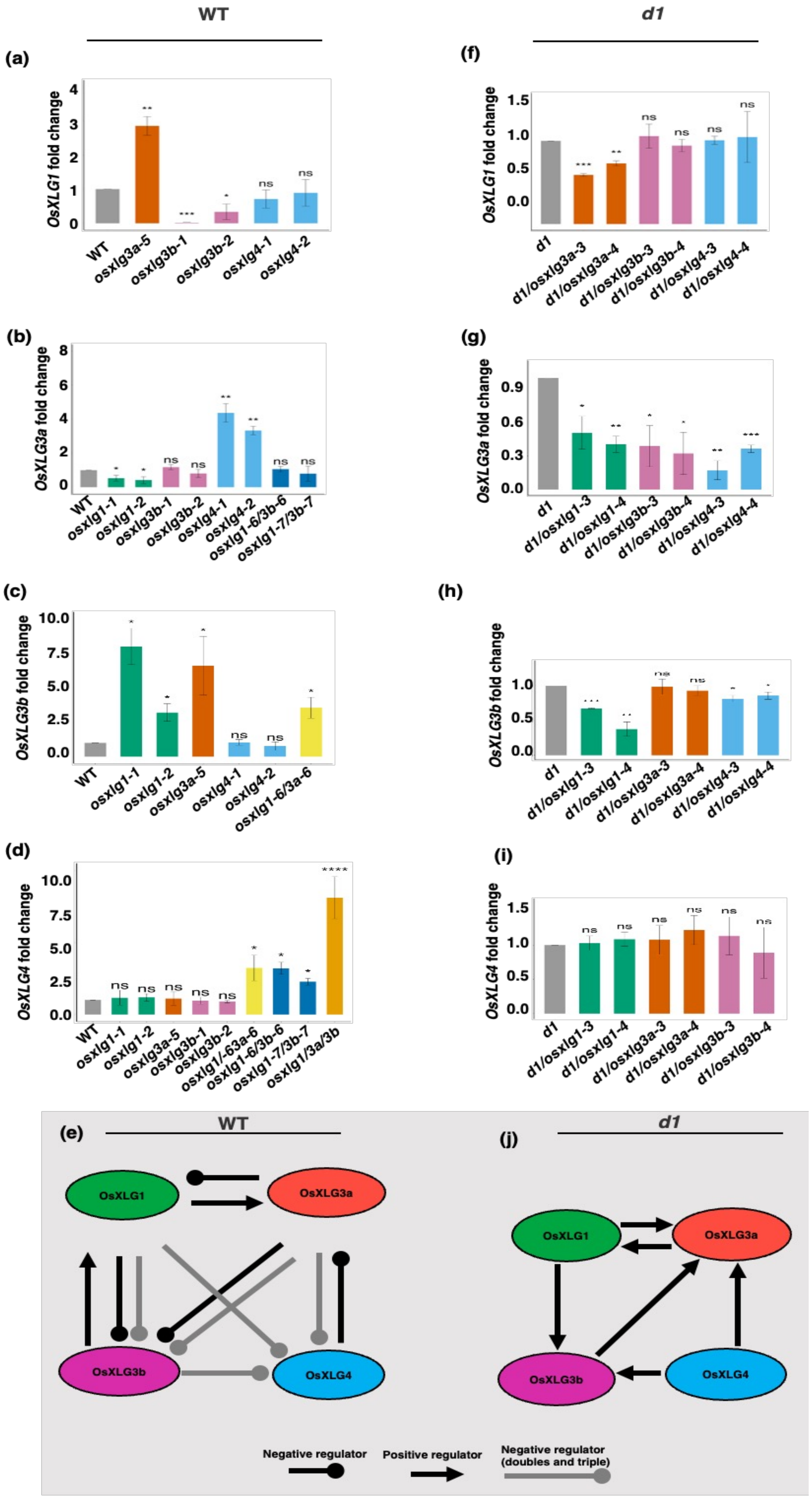


**SI, Figure 4. Analysis of *OsXLGs* expression in CRISPR mutants using qRT-PCR and proposed interactions within the rice Gα protein network, based on transcript abundance.** (a-d) *OsXLGs* expression of CRISPR mutants in WT background. (e) Regulatory relationship among OsXLGs, showcasing a feedback loop mechanism. OsXLG1 positively regulates *OsXLG3a* but negatively affects *OsXLG3b*. OsXLG3a negatively regulates *OsXLG1* and *OsXLG3b*. OsXLG3b positively influences *OsXLG1*. OsXLG4 negatively regulates *OsXLG3a*. (f-i) *OsXLGs* expression of CRISPR mutants in *d1* mutant background. (j) Regulatory relationship among OsXLGs, exhibiting a positive feedback loop. OsXLG1 positively regulates *OsXLG3a* and *OsXLG3b*, while OsXLG3a positively regulates *OsXLG1*. OsXLG3b positively influences *OsXLG3a*. OsXLG4 also plays a role in positively regulating *OsXLG3a* and *OsXLG3b* Arrows with circle ends signify negative regulation, and those with triangle ends indicate positive regulation. Statistical significance levels are indicated as follows: ns, non-significant *, $P < 0.05$; **, $P < 0.01$; ***, $P < 0.001$, ****, $P < 0.0001$.

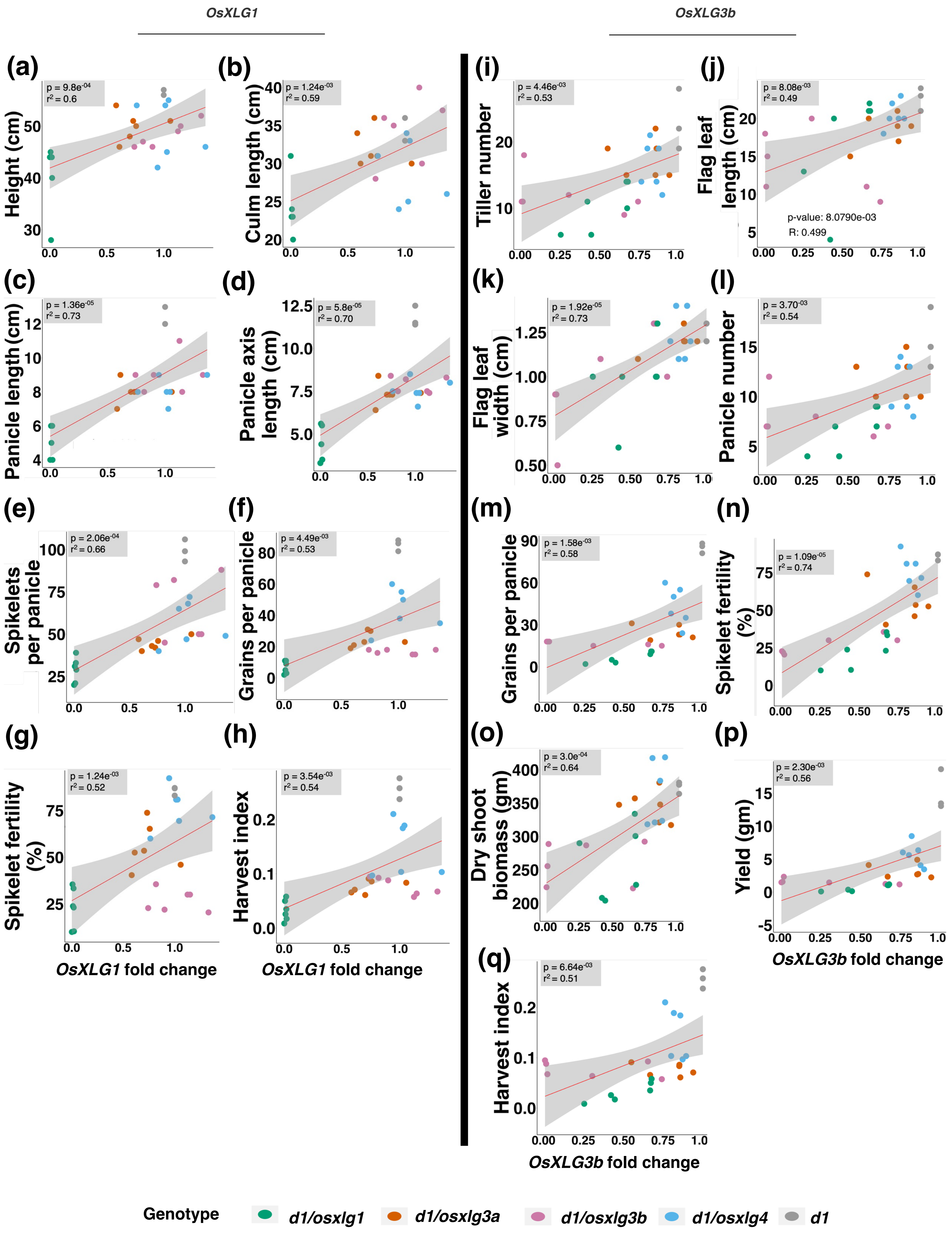


**SI, Figure 5. Correlation of *OsXLG1* and *OsXLG3a* expression with various agronomic traits in the *d1* mutant background.** Only significant correlations are shown. (a-h) *OsXLG1* expression. (i-p) *OsXLG3a* expression. The x-axis labels for *OsXLG1* fold change in panels (g-h) are applicable across panels (a-h). The x-axis labels for *OsXLG3a* fold change in panels (p-q) apply to panels (i-p).

**SI, Table 1. crRNA sequence selected for each *OsXLG* gene.**

| Chromosome region | *OsXLG* name | **PAM†** + crRNA† sequence (5'---> 3') |
|---|---|---|
| Os12g0593000 | *OsXLG1* | crRNA1- **TTTC**GTAATCTCGCGGCCGTTTATCAAGA<br>crRNA2- **TTTC**ACGCCCTTCAAGTAGTATACCAAG<br>crRNA3- **TTTC**CATAGTTCCTCAACCATTGGTGCA |
| Os11g0206700 | *OsXLG3a* | crRNA1- **TTTT**GCTTGAGGGTCGTGAACGCTTTGA<br>crRNA2- **TTTT**GGATATCATAGCGATGGGCGATCT<br>crRNA3- **TTTC**TTCCCTGCCGCAACGCGAGAATAC |
| Os06g0111400 | *OsXLG3b* | crRNA1- **TTTA**TGCCGCCGGTGCACCGGAAGCAGG<br>crRNA2- **TTTC**CATGGGCAGAGAGGTGGTAGGACA<br>crRNA3- **TTTG**CGACGCGAGGTTTTGCAGCTACTG |
| Os10g0117800 | *OsXLG4* | crRNA1- **TTTG**AGGACACCCCCTGTGTACTCCTCC<br>crRNA2- **TTTA**AGGACCTCTACTCCTCCGTCGCTG<br>crRNA3- **TTTG**TGTTCCAGACGAAGGCGCTGGAGC |

**†**PAM- proto adjacent motif, crRNA- CRISPR RNA

**SI, Table 2. Primers used in the study.**

| **Primer name** | **Sequence** (5'→ 3') |
|---|---|
| Hyg_F | AGCGAGAGCCTGACCTATTG |
| Hyg_R | GAACATCGCCTCGCTCCAG |
| Cpf1_F | TCAACGGATTCACAACAGCATTCA |
| Cpf1_R | TTTGTCTCTGATCACGTTCCATTC |
| HH_F | CTGATGAGTCCGTGAGGACG |
| HH_R | GTCCCATTCGCCATGCCGAA |
| SSTU_LB_XLGquad_F | CGATCGGGATCCGGTACCTAAT |
| SSTU_LB_XLGquad_R | CCGGATGGATCCGGTACCATCT |
| OsXLG4_F | GACTCATACGCTGACAAAGC |
| OsXLG4_R | GGAGAGGTCGTC TGTCCAC |
| OsXLG3A_F2 | GCTTTGACACGTTTTGCATGC |
| OsXLG3A_R2 | AGAAAGAATAGATACAGTTTTCT |
| OsXLG3B_F2_ST | CCATGGGCAGAGAGGTGGTA |
| OsXLG3B_R2_ST | AGCAGGAACAAAATGTCATT |
| XLG1_F5_TARGET2_3 | CATCTAGGCTGGCACCTGAC |
| XLG1_R5_TARGET2_3 | GGATGTTATCCCATCAGCGT |
| XLG1F6_TARGET1 | AAACTGATTATGACATCTC |
| XLG1R6_TARGET1 | GTCGATTTACCACATTCACT |
| OsXLG3AqPCR_F2 | GAACTCAAGGACACTCTCAAGG |
| OsXLG3AqPCR_R2 | CAGTGGACAACCATTCACAATAAG |
| OsXLG3BqPCR_F2 | AGCAGGGCCATTGAAGTATC |
| OsXLG3BqPCR_R2 | GCTACGGTCATCCAGAGTAAAC |
| OsXLG4qPCR_F2 | AGGACATCCGTGCCATAATC |
| OsXLG4qPCR_R2 | GTGCCTGATCACATCCTCAA |
| OsXLG1qPCR_F2 | GATAAACGGCCGCGAGATTA |

| | |
|---|---|
| OsXLG1qPCR_R2 | GGTAAGTCCCATCAGCATTCA |
| GAPDHqPCR_F1 | AAGCCAGCATCCTATGATCAGATT |
| GAPDHqPCR_R1 | CGTAACCCAGAATACCCTTGAGTTT |
| RGA1qPCR_F1 | CTGGGAAACAGGAGGTTGAA |
| RGA1qPCR_R1 | TAGGGCCGTAGTTCTGTAGAT |
| OsXLG3a_Biswalsite_F1 | TCCGGAAAGATTCAACCTTGT |
| OsXLG3a_Biswalsite_R1 | TGCTGTCCGGCGTGAACCCGA |
| OsXLG3b_Biswalsite_F1 | AGGTGAGGAGGAGGAGGTTA |
| OsXLG3b_Biswalsite_R1 | CGATTCCGAGCTTCTCCTCC |
| OsXLG1_Biswalsite_F1 | ATACGGCGAGGACGCCATGC |
| OsXLG1_Biswalsite_R1 | CGGCGACAGCTTGGTGCCAT |
| XLG3AOffTarget_F1 | GCCTCCGTTGTGATACCGGT |
| XLG3AOffTarget_R1 | GAGCAGGTCGGTGAGCTCGC |

**SI, Table 3. Summary of *OsXLG* mutant alleles generated by CRISPR/CpF1.**

| Background | *OsXLG* gene | No. of plants generated | No. of plants with mutation | Mutation efficiency | No. T1 lines analyzed | No. of CRISPR free and homozygous T1 lines |
|---|---|---|---|---|---|---|
| **Wild type** | *OsXLG1* | 218 | 218 | 100.00 | 30 | 18 |
| | *OsXLG3a* | 7 | 7 | 100.00 | NA | NA |
| | *OsXLG3b* | 61 | 61 | 100.00 | 30 | 11 |
| | *OsXLG4* | 53 | 118 | 44.92 | 30 | 25 |
| ***d1* mutant** | *OsXLG1* | 63 | 63 | 100.00 | 30 | 22 |
| | *OsXLG3a* | 47 | 47 | 100.00 | 30 | 21 |
| | *OsXLG3b* | 60 | 60 | 100.00 | 30 | 19 |
| | *OsXLG4* | 56 | 20 | 35.71 | 30 | 19 |

**SI, Table 4. CRISPR *OsXLG* allele description.**

| Background | *OsXLG* gene | Allele | Deletion type | Mutation type |
|---|---|---|---|---|
| **Wild type** | *OsXLG1* | *osxlg1-1* | 11bp deletion | Frameshift mutation in position 605 leading to STOP codon at position 606. |
| | | *osxlg1-2* | 11bp deletion | Frameshift mutation in position 530 leading to STOP codon. |
| | *OsXLG3a* | *osxlg3a-1* | 25bp deletion | Frameshift mutation in position 558 leading to a STOP codon at position 591. |
| | | *osxlg3a-2* | 10bp deletion | Frameshift mutation in position 564 leading to STOP codon at position 596. |
| | *OsXLG3b* | *osxlg3b-1* | 9bp deletion | Deletion of three amino acids in position 222, 223 and 224. |
| | | *osxlg3b-2* | 11bp deletion | Frameshift mutation in position 220 leading to STOP codon at position 243. |
| | *OsXLG4* | *osxlg4-1* | 8bp deletion | Frameshift mutation in position 783 leading to STOP codon at position 799. |
| | | *osxlg4-2* | 13bp deletion | Frameshift mutation in position 710 leading to STOP codon at position 723. |
| ***d1* mutant** | *OsXLG1* | *osxlg1-3* | 7bp deletion | Frameshift mutation in position 608 leading to STOP codon at position 661. |
| | | *osxlg1-4* | 11bp deletion | Frameshift mutation in position 530 leading to STOP codon. |
| | *OsXLG3a* | *osxlg3a-3* | 13bp deletion | Frameshift mutation in position 558 leading to a STOP codon at position 591. |
| | | *osxlg3a-4* | 11bp deletion | Frameshift mutation in position 564 leading to STOP codon at position 596. |
| | *OsXLG3b* | *osxlg3b-3* | 12bp deletion | Deletion of four amino acids in position 222, 223, 224 and 225. |

| | | | | |
|---|---|---|---|---|
| | | *osxlg3b-4* | 16bp deletion | Frameshift mutation in position 220 leading to STOP codon. |
| | *OsXLG4* | *osxlg4-3* | 11bp deletion | Frameshift mutation in position 783 leading to STOP codon at position 798. |
| | | *osxlg4-4* | 9bp deletion | Deletion of 3 amino acids in positions 782, 783 and 784. |

### SI, Table 5. Correlations between expression of Gα genes and *osxlg* mutant phenotypes in WT and *d1* mutant background.

| **Background →** | **WT background** | | | | | | | | | | ***d1* mutant background** | | | | | | | |
|---|---|---|---|---|---|---|---|---|---|---|---|---|---|---|---|---|---|---|
| **Gene →** | ***RGA1*** | | ***OsXLG1*** | | ***OsXLG3a*** | | ***OsXLG3b*** | | ***OsXLG4*** | | ***OsXLG1*** | | ***OsXLG3a*** | | ***OsXLG3b*** | | ***OsXLG4*** | |
| **Phenotypes** | p value | $r^2$ | p value | $r^2$ | p value | $r^2$ | p value | $r^2$ | p value | $r^2$ | p value | $r^2$ | p value | $r^2$ | p value | $r^2$ | p value | $r^2$ |
| Plant height | 5.97E-05 | 0.73 | 2.05E-01 | 0.27 | 2.06E-02 | 0.47 | 3.88E-02 | 0.42 | 4.18E-03 | 0.56 | 9.80E-04 | 0.60 | 2.52E-01 | 0.23 | 6.16E-02 | 0.37 | 8.77E-01 | 0.03 |
| Tiller number | 1.75E-02 | 0.48 | 1.68E-01 | 0.29 | 5.65E-01 | 0.12 | 6.40E-03 | 0.54 | 3.89E-01 | 0.18 | 2.13E-02 | 0.44 | 5.20E-01 | 0.13 | 4.46E-03 | 0.53 | 9.77E-01 | 0.01 |
| Flag leaf length | 2.03E-01 | 0.27 | 9.06E-01 | 0.03 | 1.35E-04 | 0.70 | 2.35E-01 | 0.25 | 9.31E-04 | 0.63 | 5.18E-01 | 0.13 | 4.25E-01 | 0.16 | 8.08E-03 | 0.50 | 8.45E-02 | 0.34 |
| Flag leaf width | 6.42E-02 | 0.38 | 2.69E-01 | 0.24 | 2.05E-01 | 0.27 | 9.12E-03 | 0.52 | 2.35E-01 | 0.25 | 3.84E-01 | 0.17 | 6.80E-01 | 0.08 | 1.92E-05 | 0.73 | 3.57E-01 | 0.18 |
| Culm length | 2.12E-04 | 0.69 | 1.70E-01 | 0.29 | 3.75E-01 | 0.19 | 8.82E-02 | 0.36 | 1.18E-01 | 0.33 | 1.24E-03 | 0.59 | 6.17E-01 | 0.10 | 1.00E-01 | 0.00 | 1.28E-01 | 0.30 |
| Days-to-heading | 8.93E-01 | 0.03 | 1.59E-01 | 0.30 | 2.42E-01 | 0.25 | 6.91E-01 | 0.09 | 3.13E-01 | 0.22 | 4.50E-01 | 0.15 | 1.90E-01 | 0.26 | 2.45E-02 | 0.43 | 2.71E-01 | 0.22 |
| Panicle length | 1.04E-02 | 0.51 | 3.10E-02 | 0.44 | 2.85E-01 | 0.23 | 9.25E-01 | 0.02 | 1.45E-01 | 0.31 | 1.37E-05 | 0.73 | 2.77E-02 | 0.42 | 6.48E-02 | 0.36 | 7.31E-01 | 0.07 |
| Panicle exsertion length | 8.30E-03 | 0.53 | 1.67E-01 | 0.29 | 1.15E-01 | 0.33 | 6.59E-01 | 0.10 | 6.49E-02 | 0.38 | 3.80E-01 | 0.18 | 7.50E-02 | 0.35 | 2.05E-02 | 0.44 | 1.64E-01 | 0.28 |
| Panicle axis length | 8.05E-02 | 0.36 | 1.30E-02 | 0.50 | 8.18E-01 | 0.05 | 4.92E-01 | 0.15 | 5.17E-01 | 0.14 | 5.80E-05 | 0.70 | 3.04E-02 | 0.42 | 5.50E-02 | 0.37 | 9.51E-01 | 0.01 |
| Panicle number | 7.31E-02 | 0.37 | 1.58E-02 | 0.49 | 4.45E-01 | 0.16 | 1.42E-02 | 0.49 | 5.37E-01 | 0.13 | 2.39E-02 | 0.43 | 4.90E-01 | 0.14 | 3.70E-03 | 0.54 | 1.00E-01 | 0.01 |
| Grain per panicle | 2.12E-02 | 0.47 | 8.76E-03 | 0.52 | 4.29E-01 | 0.17 | 3.26E-01 | 0.21 | 5.29E-01 | 0.14 | 4.85E-08 | 0.53 | 1.28E-02 | 0.47 | 1.58E-03 | 0.58 | 9.57E-01 | 0.01 |
| Spikelet per panicle | 4.31E-02 | 0.42 | 2.48E-02 | 0.46 | 4.94E-01 | 0.15 | 1.75E-01 | 0.29 | 5.29E-01 | 0.14 | 2.06E-04 | 0.66 | 2.30E-02 | 0.44 | 7.48E-01 | 0.07 | 1.58E-01 | 0.28 |
| Spikelet fertility | 9.70E-02 | 0.35 | 1.63E-02 | 0.49 | 2.76E-01 | 0.23 | 7.21E-01 | 0.08 | 3.98E-01 | 0.18 | 5.54E-03 | 0.52 | 4.78E-01 | 0.14 | 1.10E-05 | 0.74 | 5.40E-01 | 0.12 |
| Grain length | 2.34E-02 | 0.46 | 3.48E-01 | 0.20 | 2.34E-06 | 0.80 | 5.00E-04 | 0.66 | 1.79E-05 | 0.76 | 7.28E-01 | 0.07 | 1.53E-02 | 0.46 | 3.95E-01 | 0.17 | 4.19E-01 | 0.16 |
| Grain width | 3.56E-01 | 0.20 | 9.02E-01 | 0.03 | 1.48E-02 | 0.49 | 2.08E-01 | 0.27 | 4.37E-02 | 0.42 | 5.00E-01 | 0.15 | 7.53E-02 | 0.35 | 9.36E-01 | 0.02 | 4.72E-01 | 0.14 |
| 100-Grain weight | 7.77E-02 | 0.37 | 2.76E-01 | 0.23 | 6.61E-05 | 0.72 | 4.22E-02 | 0.42 | 1.34E-03 | 0.62 | 3.68E-01 | 0.18 | 2.23E-01 | 0.24 | 6.69E-01 | 0.09 | 4.10E-02 | 0.40 |
| Biomass | 3.05E-01 | 0.22 | 1.02E-01 | 0.34 | 1.40E-01 | 0.31 | 1.01E-01 | 0.34 | 3.92E-01 | 0.18 | 1.02E-02 | 0.49 | 7.69E-01 | 0.06 | 3.00E-04 | 0.64 | 3.00E-01 | 0.21 |
| Yield | 3.28E-01 | 0.21 | 4.70E-04 | 0.66 | 8.71E-01 | 0.04 | 9.16E-01 | 0.02 | 7.94E-01 | 0.06 | 2.33E-02 | 0.44 | 2.70E-03 | 0.55 | 2.30E-03 | 0.56 | 7.92E-01 | 0.05 |
| Harvest index | 9.88E-03 | 0.52 | 1.26E-01 | 0.32 | 6.97E-01 | 0.08 | 9.44E-02 | 0.35 | 9.81E-01 | 0.01 | 3.54E-03 | 0.54 | 7.69E-01 | 0.06 | 3.00E-04 | 0.64 | 3.00E-01 | 0.21 |

**SI, Table 6. Genetic interaction between *RGA1* and *OsXLGs*.** The average phenotype values for the CRISPR alleles of each *OsXLG* gene and *d1* allele were calculated. The individual effect of each Gα mutation was calculated by subtracting the averaged phenotype values from the Nipponbare wild type values. The “expected effect” of the combined Gα mutations was then calculated by adding the individual effects (*d1* + *osxlg*). The “actual effect” was calculated by subtracting the average phenotype value of the plants with both mutations (*d1/osxlg*) to Nipponbare wild type. The “expected effect” was compared with the “actual effect”. Based on these comparisons, we categorized the interactions as synergistic (greater than the additive effect), additive (equal to the expected effect), partially redundant (less than the additive effect but still significant), and we identified suppressive, enhancing, or antagonistic effects of Gα mutations. In the expected effect, phenotypic categories include "down" or "up" for a phenotype lower or higher than the wild type (WT) in the single gene mutation effect, with "ns" indicating no significance compared to WT. In the actual effect, "down" represents a double mutant phenotype lower than WT but not lower than either single gene mutation effect, while "down/down" indicates a double mutant with a phenotype lower than either the WT or single mutation effect. Conversely, "up" signifies a double mutant phenotype higher than WT but not lower than either single gene mutation, and "up/up" indicates a double mutant with a phenotype higher than either the WT or single mutation effect. “No change” indicates that the actual effect is the same as the combined effect. (-, -) denotes significance of either the *d1* or *osxlg* single mutant phenotype; (ns, -) indicates no significance for the *d1* but significance of *osxlg* single mutant phenotype; (-,ns) signifies significance to *d1* but not significant to *osxlg* single mutant phenotype; and (ns,ns) denotes no significance to either *d1* or *osxlg* single mutant phenotype.
For example, regarding plant height of *d1* and *osxlg1* interaction: the expected effect of the *d1* mutant phenotypeis "down" (lower than WT), while the expected effect of the *osxlg1* mutant phenotype is nonsignificant (ns). The actual effect of the *d1/osxlg1* double mutant phenotype is "down/down," indicating further reduction compared to the combined effect of *d1* and *osxlg1* single mutant phenotype. (-, -) suggests that the actual effect of the *d1/osxlg1* mutant phenotype is significant to either *d1* or *osxlg1* single mutant phenotype effect.

| **Phenotype** | ***OsXLG1*** | | ***OsXLG3a*** | | ***OsXLG3b*** | | ***OsXLG4*** | |
|---|---|---|---|---|---|---|---|---|
| Interaction → | (*d1* + *osxlg1*) = expected effect | (*d1 osxlg1*) = actual effect | (*d1* + *osxlg3a*) = expected effect | (*d1 osxlg3a*) = actual effect | (*d1* + *osxlg3b*) = expected effect | (*d1 osxlg3b*) = actual effect | (*d1* + *osxlg4*) = expected effect | (*d1 osxlg4*) = actual effect |
| Plant height | 34.8 cm + (-0.1 cm) = **34.7 cm** **down + ns** | **51.3 cm** **(- , -)** | 34.8 cm + 16.6 cm = **51.4 cm** **down +down** | **41.1 cm** **(- , -)** | 34.8 cm + 16 cm = **50.8 cm** **down + down** | **43.9 cm** **(-,-)** | 34.8 cm + 20.5 cm = **55.3 cm** **down +down** | **40.9 cm** **(-,-)** |
| | There is an enhancing effect of *osxlg1* on *d1.* | | *d1* and *osxlg3a are* partially redundant. | | *d1* and *osxlg3b are* partially redundant. | | *d1* and *osxlg4 are* partially redundant. | |
| Flag leaf length | 9 cm + (-1.6 cm) = **7.4 cm** **down + ns** | **15.2 cm** **down/down** **(-,-)** | 9 cm + (-2 cm) = **7 cm** **down + ns** | **13.6 cm** **down/down** **(-,-)** | 9 cm + (-2.5 cm) = **6.5 cm** **down + ns** | **17.3 cm** **down/down** **(-,-)** | 9 cm + 6.3 cm = **15.3 cm** **down + down** | **10.2 cm** **down** **(ns, -)** |

| | | | | | | | | |
|---|---|---|---|---|---|---|---|---|
| | There is an enhancing effect of *osxlg1* on *d1.* | | There is an enhancing effect of *osxlg3a* on *d1.* | | There is an enhancing effect of *osxlg3b* on *d1.* | | *d1* is epistatic to *osxlg4.* | |
| Flag leaf width | (-.24 cm) + (-.02 cm)<br>= **-0.26 cm**<br>**ns + ns** | **-0.08 cm**<br>**up**<br>**(ns, ns)** | (-.24 cm) + (0 cm)<br>= **-0.24 cm**<br>**ns + ns** | **-0.24 cm**<br>**No change**<br>**(ns, ns)** | (-.24 cm) + (-.04 cm)<br>= **-0.28 cm**<br>**ns+ ns** | **-0.05 cm**<br>**up**<br>**(ns, ns)** | (-.24 cm) +(-.04 cm) =<br>**-0.28 cm**<br>**ns + ns** | **-0.22 cm**<br>**up**<br>**(ns, ns)** |
| | *d1* is independent. | | *d1* is independent. | | *d1* is independent. | | *d1* is independent. | |
| Culm length | 25.8 cm+ 10.3 cm<br>= **36.1 cm**<br>**down + down** | **36.1 cm**<br>**no change**<br>**(-,-)** | 25.8 cm+ 18.6 cm<br>= **44.4 cm**<br>**down + down** | **26.7 cm**<br>**down**<br>**(-,-)** | 25.8 cm+ 18.5 cm<br>= **44.3 cm**<br>**down + down** | **26.6 cm**<br>**down**<br>**(-,-)** | 25.8 cm+ 14.2cm<br>= **40 cm**<br>**down + down** | **30.7 cm**<br>**down**<br>**(-,-)** |
| | *d1* is additive with *osxlg1.* | | *d1* and *osxlg3a are* partially redundant. | | *d1* and *osxlg3b are* partially redundant. | | *d1* and *osxlg4 are* partially redundant. | |
| Tiller number | 0.8 + 2.7 = **3.5**<br>**ns + ns** | **11.5**<br>**down**<br>**(- , -)** | 0.8 + 0.2 = **1**<br>**ns + ns** | **6.3**<br>**down**<br>**(-, ns)** | 0.8+ (-17.5) = **-16.7**<br>**ns + up** | **11.4**<br>**down**<br>**(-,-)** | 0.8 + (-2.9) = **-2.1**<br>**ns + ns** | **6**<br>**down**<br>**(- , -)** |
| | Redundant. | | Redundant. | | *d1* antagonizes *osxlg3b.* | | Redundant. | |
| Days to heading | 25.2 days + 35.9 days = **61.1 days**<br>**down + down** | **28.6 days**<br>**down**<br>**(ns, -)** | 25.2 days+ 43.8 days = **69 days**<br>**down + down** | **31.1 days**<br>**down**<br>**(ns, -)** | 25.2 days + 61.7 days= **86.9 days**<br>**down + down** | **32.3 days**<br>**down**<br>**(ns, -)** | 25.2 days+ 62.7 days<br>= **87.9 days**<br>**down + down** | **28.1 days**<br>**down**<br>**(ns, -)** |
| | *d1* is epistatic to *osxlg1.* | | *d1* is epistatic to *osxlg3a.* | | *d1* is epistatic to *osxlg3b.* | | *d1* is epistatic to *osxlg4.* | |
| Panicle length | 9.6 cm + 7.5 cm<br>= **17.1 cm**<br>**down + down** | **16.4 cm**<br>**down**<br>**(- , -)** | 9.6 cm + 8.4 cm<br>= **18 cm**<br>**down + down** | **13.8 cm**<br>**down**<br>**(- , -)** | 9.6 cm + 5.8 cm<br>= **15.4 cm**<br>**down + down** | **13 cm**<br>**down**<br>**(- , -)** | 9.6 cm + 6.5 cm<br>= **16.1 cm**<br>**down + down** | **14.1 cm**<br>**down**<br>**(- , -)** |
| | *d1* and *osxlg1* are partially redundant. | | *d1* and *osxlg3a* are partially redundant. | | *d1* and *osxlg3b* are partially redundant. | | *d1* and *osxlg4* are partially redundant. | |
| Panicle exsertion length | 4.7 cm + 3.6 cm<br>= **8.3 cm**<br>**down + down** | **4.73 cm**<br>**down**<br>**(ns, -)** | 4.7 cm + 4.8 cm<br>= **9.5 cm**<br>**down + down** | **4.66 cm**<br>**down**<br>**(ns, ns)** | 4.7 cm + 4.2 cm<br>= **8.9 cm**<br>**down + down** | **4.67 cm**<br>**down**<br>**(ns, ns)** | 4.7 cm + 4.4 cm<br>= **9.1 cm**<br>**down + down** | **4.77 cm**<br>**down**<br>**(ns, ns)** |
| | *d1* is epistatic to *osxlg1.* | | *d1* is epistatic to *osxlg3a.* | | *d1* is epistatic to *osxlg3b.* | | *d1* is epistatic to *osxlg4.* | |
| Panicle axis length | 4.88 cm+ 3.9 cm<br>= **8.8 cm**<br>**down + ns** | **11.7 cm**<br>**down/down**<br>**(-,-)** | 4.88 cm + 3.6 cm<br>= **8.5 cm**<br>**down + down** | **9.1 cm**<br>**down/down**<br>**(-, -)** | 4.88 cm + 1.6 cm<br>= **6.5 cm**<br>**down + down** | **8.6 cm**<br>**down/down**<br>**(-,-)** | 4.88 cm+ 2.1 cm<br>= **7.01 cm**<br>**down + down** | **9.4 cm**<br>**down/down**<br>**(-,-)** |
| | There is an enhancing effect of *osxlg1* on *d1.* | | *d1* and *osxlg3a* are synergistic. | | *d1* and *osxlg3b* are synergistic. | | *d1* and *osxlg4* are synergistic. | |
| Panicle number | -1.4 + (-3.2) = **-4.6**<br>**ns + ns** | **6.1**<br>**down**<br>**(-,-)** | -1.4 + 4 = **2.6**<br>**ns + ns** | **2.3**<br>**down**<br>**(-, ns)** | -1.4 + (-13.5) = **-**<br>**14.9**<br>**ns + up** | **6**<br>**down**<br>**(-,-)** | -1.4 + (-3.3) = **-4.7**<br>**ns + ns** | **2.3**<br>**down**<br>**(-,-)** |
| | Redundant. | | Redundant. | | *d1* antagonizes *osxlg3b.* | | Redundant. | |
| Spikelets per panicle | -15.6 + (-3.8) = **-19.4**<br>**up + up** | **52.6**<br>**down**<br>**(-,-)** | -15.6 + 16.6 = **1**<br>**up + down** | **39.5**<br>**down/down**<br>**(-,-)** | -15.6 + 12.1 = **-3.5**<br>**up + down** | **17**<br>**down/down**<br>**(-,-)** | -15.6 + (-19.2) = **-34.8**<br>**up + up** | **24.7**<br>**down**<br>**(-,-)** |
| | *d1* and *osxlg1* are mutually antagonistic. | | *osxlg3a* is epistatic to *d1* and there is an enhancing effect of *d1* on *osxlg3a.* | | *osxlg3b* is epistatic to *d1* and there is an enhancing effect of *d1* on *osxlg3b.* | | *d1* and *osxlg4* are mutually antagonistic. | |

| | | | | | | | | |
|---|---|---|---|---|---|---|---|---|
| Grains per panicle | -17.6 + (-12.8)<br>= **-4.8**<br>**up + up** | **63.7**<br>**down/down**<br>**(-,-)** | -17.6 + 31.6<br>= **14**<br>**up + down** | **45.1**<br>**down/down**<br>**(-,-)** | -17.6 + 19.8<br>= **2.2**<br>**up + down** | **52.5**<br>**down/down**<br>**(-,-)** | -17.6 + (-12.5)<br>= **-30.1**<br>**up + up** | **25.5**<br>**down**<br>**(-,-)** |
| | *d1* and *osxlg1* are mutually antagonistic. | | *osxlg3a* is epistatic to *d1* and there is an enhancing effect of *d1* on *osxlg3a.* | | *osxlg3b* is epistatic to *d1* and there is an enhancing effect of *d1* on *osxlg3b.* | | *d1* and *osxlg4* are mutually antagonistic. | |
| Spikelet fertility | -8.54 % + 21.8 %<br>= **-4.06 %**<br>**ns + down** | **64 %**<br>**down/down**<br>**(-,-)** | -4.6 % + 26.5 %<br>= **22 %**<br>**ns + down** | **26.8 %**<br>**down/down**<br>**(-, ns)** | -4.6 % + 14 %<br>= **9.4 %**<br>**ns + down** | **56.2%**<br>**down/down**<br>**(-,-)** | -4.6 % + 3.2 %<br>= **-1.4 %**<br>**ns + ns** | **9.2 %**<br>**down/down**<br>**(-,-)** |
| | There is an enhancing effect of *d1* on *osxlg1.* | | *osxlg3a* is independent of *d1.* | | there is an enhancing effect of *d1* on *osxlg3b.* | | Redundant. | |
| Grain length | 1.6 mm + (-.02 mm)<br>= **1.6 mm**<br>**down + ns** | **2.51 mm**<br>**down/down**<br>**(-,-)** | 1.6 mm + (-.5 mm) =<br>**1.12 mm**<br>**down + ns** | **2.89 mm**<br>**down/down**<br>**(-,-)** | 1.6 mm + .38 mm<br>= **1.98 mm**<br>**down + ns** | **2.78 mm**<br>**down/down**<br>**(-,-)** | 1.6 mm + 1.04 mm =<br>**2.64 mm**<br>**down + down** | **2.29 mm**<br>**down**<br>**(-,-)** |
| | There is an enhancing effect of *osxlg1* on *d1.* | | There is an enhancing effect of *osxlg3a* on *d1.* | | There is an enhancing effect of *osxlg3b* on *d1.* | | *d1* and *osxlg4* are partially redundant. | |
| Grain width | -0.14 mm + (-0.15 mm) = **-0.29 mm**<br>**ns + ns** | **0.02 mm**<br>**(ns, -)** | -0.14 mm + (-0.34 mm) = **-0.48 mm**<br>**ns + ns** | **0.58 mm**<br>**down**<br>**(-,-)** | -0.14 mm + (-0.2 mm) = **-0.34 mm**<br>**ns + ns** | **0.25 mm**<br>**down**<br>**(-, ns)** | -0.14 mm + 0.1 mm =<br>**-0.05 mm**<br>**ns + ns** | **-0.11 mm**<br>**(ns, ns)** |
| | No effect. | | Redundant. | | Redundant. | | No effect. | |
| 100- grain weight | 1.3 g + 0.01 g<br>= **1.32 g**<br>**down + ns** | **1.61 g**<br>**down/down**<br>**(-,-)** | 1.3 g + 0.02 g<br>= **1.3 g**<br>**down + ns** | **1.7 g**<br>**down/down**<br>**(-,-)** | 1.3 g + 0.2 g<br>= **1.5 g**<br>**down + down** | **1.6 g**<br>**down/down**<br>**(-,-)** | 1.3 g + 0.5 g<br>= **1.77 g**<br>**down + down** | **1.53 g**<br>**down**<br>**(-,-)** |
| | There is an enhancing effect of *osxlg1* on *d1.* | | There is an enhancing effect of *osxlg3a* on *d1.* | | *d1* and *osxlg3b* are synergistic. | | *d1* and *osxlg4* are partially redundant. | |
| Yield | 9 g + 5 g<br>= **14 g**<br>**down + down** | **24.3 g**<br>**down/down**<br>**(-,-)** | 9 g + 16 g<br>= **25 g**<br>**down + down** | **22 g**<br>**down**<br>**(-,-)** | 9 g + (-7 g)<br>= **2 g**<br>**down + up** | **23.2 g**<br>**down/down**<br>**(-,-)** | 9 g + (-4 g)<br>= **4.9 g**<br>**down + ns** | **18.9 g**<br>**down/down**<br>**(-,-)** |
| | *d1* and *osxlg1* are synergistic. | | *d1* and *osxlg3a* are partially redundant | | *d1* is epistatic to *osxlg3b* and there is an enhancing effect of *osxlg3b* on *d1.* | | There is an enhancing effect of *osxlg4* on *d1.* | |
| Biomass | 186 g + (-23 g)<br>= **163 g**<br>**down + ns** | **310 g**<br>**down/down**<br>**(-,-)** | 186 g + 71 g<br>= **257 g**<br>**down + down** | **242 g**<br>**down**<br>**(-,-)** | 186 g + (-205 g)<br>= **-18.7 g**<br>**down + up** | **321 g**<br>**down/down**<br>**(-,-)** | 186 g + 95 g<br>= **280 g**<br>**down + down** | **221 g**<br>**down**<br>**(-,-)** |
| | There is an enhancing effect of *osxlg1* on *d1.* | | *d1* and *osxlg3a* are partially redundant. | | *d1* is epistatic to *osxlg3b* and there is an enhancing effect of *osxlg3b on d1.* | | *d1* and *osxlg4* are partially redundant. | |
| Harvest index | 0.05 + (-0.13) = **-.08**<br>**ns + up** | **0.27**<br>**down**<br>**(-,-)** | 0.05 + 0.12 = **0.16**<br>**ns + down** | **0.22**<br>**down/down**<br>**(-,-)** | 0.05 + 0.15 = **0.2**<br>**ns + down** | **0.2**<br>**No change**<br>**(-,-)** | 0.05 + (-0.06)<br>= **0.005**<br>**ns + up** | **.15**<br>**down**<br>**(-,-)** |
| | *d1* suppresses *osxlg1.* | | *d1* and *osxlg3a* are synergistic. | | *d1* is additive with *osxlg3b.* | | *d1* suppresses *osxlg4.* | |

**SI, Table 7. Genetic interaction between *OsXLGs*.** The average phenotype values for the alleles of each *OsXLG* gene were calculated. The individual effect of each osxlg mutation was calculated by subtracting the averaged phenotype values from Nipponbare wild type values. The "expected effect" of the combined *osxlg* mutations was then calculated by adding the individual effects (*osxlg* + *osxlg*). The "actual effect" was calculated by subtracting the average phenotype value of the plants with both mutations (*osxlg/osxlg*) to Nipponbare wild type. The "expected effect" was compared with the "actual effect". Based on these comparisons, we categorized the interactions as synergistic (greater than the additive effect), additive (equal to the expected effect), partially redundant (less than the additive effect but still significant), and identified suppressive, enhancing, or antagonistic effects of *OsXLG* mutations. In the expected effect, phenotypic categories include "down" or "up" for a phenotype lower or higher than the wild type (WT) in the single gene mutation effect, with "ns" indicating no significance compared to WT. In the actual effect, "down" represents a double mutant phenotype lower than WT but not lower than either single gene mutation effect, while "down/down" indicates a double mutant with a phenotype lower than either the WT or single mutation effect. Conversely, "up" signifies a double mutant phenotype higher than WT but not lower than either single gene mutation, and "up/up" indicates a double mutant with a phenotype higher than either the WT or single mutation effect. "ns" denotes no significance compared to WT.

For example, plant height of *osxlg1* and *osxlg3a* interaction: the expected effect of the *osxlg1* mutant phenotype is nonsignificant (0), while the expected effect of the *osxlg3a* mutant phenotype is "down" (lower than WT). The actual effect of the *osxlg1/osxlg3a* double mutant phenotype is "down," indicating no further reduction compared to the *osxlg3a* single mutant phenotype.

| **Phenotype** | ***OsXLG1/OsXLG3a*** | | ***OsXLG1/OsXLG3b*** | | ***OsXLG1/OsXLG3a/ OsXLG3b*** | |
|---|---|---|---|---|---|---|
| Interaction → | (*osxlg1 + osxlg3a*) = expected effect | (*osxlg1 osxlg3a*) =<br>actual effect | (*osxlg1 + osxlg3b*) = expected effect | (*osxlg1 osxlg3b*) = actual effect | (*osxlg1 + osxlg3a + osxlg3b*) = expected effect | (*osxlg1 osxlg3a osxlg3b*) = actual effect |
| Plant height | (-0.1 cm) + 16.6 cm<br>= **16.5 cm**<br>**ns + down** | **7.6 cm**<br>**down** | (-0.1 cm) + 16 cm<br>= **15.9 cm**<br>**ns + down** | **11.1 cm**<br>**down** | (-0.1 cm) + 16.6 cm + 16 cm<br>= **32.5 cm**<br>**ns + down + down** | **11.1 cm**<br>**down** |
| | *osxlg1* suppresses *osxlg3a.* | | *osxlg1* suppresses *osxlg3b.* | | *osxlg1* suppresses *osxlg3a* and *osxlg3b*. | |
| Flag leaf length | (-1.6 cm) + (-2 cm)<br>= **-3.6 cm**<br>**ns + ns** | **-1.2 cm**<br>**ns** | (-1.6 cm) + (-2.5 cm)<br>= **-4.1 cm**<br>**ns + ns** | **-3.1 cm**<br>**ns** | (-1.6 cm) + (-2 cm) + (-2.5 cm)<br>= **-6.1 cm**<br>**ns + ns + ns** | **-2.33 cm**<br>**0** |
| | No effect. | | No effect. | | No effect. | |
| Flag leaf width | (-0.02 cm) + 0 cm<br>= **-0.02 cm**<br>**ns + ns** | **0.06 cm**<br>**ns** | (-0.02 cm) + (-0.04 cm) =<br>**-0.06 cm**<br>**ns + ns** | **-0.1 cm**<br>**ns** | (-0.02 cm) + 0 cm + (-0.04 cm)<br>= **-0.06 cm**<br>**ns + ns + ns** | **-0.04 cm**<br>**ns** |

| | | | | | | |
|---|---|---|---|---|---|---|
| | No effect. | | No effect. | | No effect. | |
| Culm length | 10.3 cm + 18.6 cm<br>= **28.9 cm**<br>**ns + down** | **8.8 cm**<br>**down** | 10.3 cm + 18.5 cm<br>= **28.8 cm**<br>**ns + down** | **17.2 cm**<br>**down** | 10.3 cm+18.6 cm+ 18.5 cm<br>= **47.35 cm**<br>**ns + down + down** | **13.4 cm**<br>**down** |
| | *osxlg1* suppresses *osxlg3a.* | | *osxlg1* suppresses *osxlg3b.* | | *osxlg1* suppresses *osxlg3a* and *osxlg3b.* | |
| Tiller number | 2.7 + 0.2 = **2.9**<br>**ns + ns** | **10.2**<br>**down** | 2.7 + (-17.5) = **-14.8**<br>**ns + up** | **-11.7**<br>**up** | 2.7 + 0.2 + (-17.5) = **-14.6**<br>**ns + ns + up** | **-.3**<br>**up** |
| | Redundant. | | *osxlg1* suppresses *osxlg3b.* | | *osxlg3a* and *osxlg1* suppresses *osxlg3b.* | |
| Days to heading | 35.9 days + 43.8 days =<br>**79.7 days**<br>**down + down** | **54.4 days**<br>**down** | 35.9 days + 61.7 days =<br>**97.6 days**<br>**down + down** | **21.7 days**<br>**down** | 35.9 days + 43.8 days + 61.7<br>days = **141.4 days**<br>**down + down + down** | **15.2 days**<br>**down** |
| | *osxlg1* and *osxlg3a* are partially redundant. | | *osxlg1* and *osxlg3b* are partially redundant. | | *osxlg1, osxlg3a* and *osxlg3b* are partially redundant. | |
| Panicle length | 7.5 cm+ 8.4 cm<br>= **14.2 cm**<br>**down + down** | **8.2 cm**<br>**down** | 7.5 cm+ 5.8 cm<br>= **13.3 cm**<br>**down + down** | **8.9 cm**<br>**down** | 7.5 cm + 8.4 cm + 5.8 cm<br>= **21.7 cm**<br>**down + down + down** | **10.8 cm**<br>**down** |
| | *osxlg1* and *osxlg3a* are partially redundant. | | *osxlg1* and *osxlg3b* are partially redundant. | | *osxlg1, osxlg3a* and *osxlg3b* are partially redundant. | |
| Panicle exsertion length | 3.6 cm + 4.8 cm<br>= **8.4 cm**<br>**down + down** | **4.3 cm**<br>**down** | 3.6 cm + 4.2 cm<br>= **7.7 cm**<br>**down + down** | **4.27 cm**<br>**down** | 3.6 cm + 4.8 cm + 4.2 cm<br>= **12.54 cm**<br>**down + down + down** | **4.1 cm**<br>**down** |
| | *osxlg1* and *osxlg3a* are partially redundant. | | *osxlg1* and *osxlg3b* are partially redundant. | | *osxlg1, osxlg3a* and *osxlg3b* are partially redundant. | |
| Panicle axis length | 3.9 cm + 3.6 cm<br>= **7.53 cm**<br>**ns + down** | **3.9 cm**<br>**down** | 3.9 cm + 1.61 cm<br>= **5.54 cm**<br>**ns + down** | **4.53 cm**<br>**down** | 3.9 cm + 3.6 cm + 1.61 cm<br>= **9.14 cm**<br>**ns + down + down** | **6.65 cm**<br>**down** |
| | *osxlg1* suppresses *osxlg3a.* | | *osxlg1* suppresses *osxlg3b.* | | *osxlg1* suppresses *osxlg3a* and *osxlg3b.* | |
| Panicle number | (-3.2) + 4 = **0.78**<br>**ns + ns** | **5.8**<br>**down** | (-3.2) + (-13.5) = **-16.7**<br>**ns + up** | **-9.9**<br>**up** | (-3.2) +4+ (-13.5) = **-12.7**<br>**ns + ns + up** | **-4.15**<br>ns |
| | *osxlg1* and *osxlg3a* are redundant. | | *osxlg1* suppresses *osxlg3b.* | | *osxlg3a* and *osxlg1* suppress *osxlg3b.* | |
| Spikelets per panicle | (-3.8) + 16.6 = **12.8**<br>**up + down** | **8.6**<br>**ns** | (-3.8) + 12.1 = **8.3**<br>**up + down** | **9.9**<br>**down/down** | (-3.8) + 16.6 + 12.1 = **25**<br>**up + down + down** | **8**<br>**down** |
| | *osxlg1 is* epistatic to *osxlg3a* with suppressing effect of *osxlg1* to *osxlg3a.* | | *osxlg1 is* epistatic to *osxlg3b* with suppressing effect of *osxlg1* to *osxlg3b* | | *osxlg1* is epistatic to *osxlg3a* and *osxlg3b* with suppressing effect of *osxlg1* to *osxlg3a* and *osxlg3b.* | |
| Grains per panicle | 12.8 + 31.6 = **44.4**<br>**down + down** | **17.8**<br>**down** | 12.8 + 19.8 = **32.6**<br>**down + down** | **31.3**<br>**down** | 12.8 31.6 + 19.8 = **64.2**<br>**down + down + down** | **42.4**<br>**down** |
| | *osxlg1* and *osxlg3a* are partially redundant. | | *osxlg1* and *osxlg3b* are partially redundant. | | *osxlg1, osxlg3a* and *osxlg3b* are partially redundant. | |
| Spikelet fertility | (-4.06 %) + 26.54 %<br>= **22.5 %**<br>**ns + down** | **13.54 %**<br>**ns** | (-4.06 %) + 14 %<br>= **9.9 %**<br>**ns + down** | **31.1 %**<br>**down** | (-4.06 %) + 26.54 % + 14 %<br>= **36.47 %**<br>**ns + down + down** | **47.6 %**<br>**down** |
| | *osxlg1* suppresses *osxlg3a.* | | There is enhancing effect of *osxlg1* to *osxlg3b.* | | There is enhancing effect of *osxlg1* to *osxlg3a* and *osxlg3b.* | |

| | | | | | | |
|---|---|---|---|---|---|---|
| Grain length | (-0.02 mm) + (-0.48 mm)<br>= **-0.5 mm**<br>**ns + ns** | **-0.58 mm**<br>**up** | (-0.02 mm) + 0.38 mm<br>= **0.36 mm**<br>**ns + down** | **0.52 mm**<br>**down** | (-0.02 mm) + -0.48 mm+ 0.38 mm<br>= **-0.12 mm**<br>**ns + ns + down** | **-0.28 mm**<br>**up** |
| | *osxlg1* and *osxlg3a* are redundant. | | There is an enhancing effect of *osxlg1* to *osxlg3b.* | | There is an enhancing effect of *osxlg1* and *osxlg3a* to *osxlg3b.* | |
| Grain width | (-0.15 mm) + (-0.34 mm) =<br>**-0.49 mm**<br>**ns + ns** | **-0.2 mm**<br>**ns** | (-0.15 mm) + (-0.2 mm) =<br>**-0.35 mm**<br>**ns + ns** | **-0.08 mm**<br>**ns** | (-0.15 mm) + (-0.34 mm) + (-0.2 mm) = **-0.69 mm**<br>**ns + ns + ns** | **0.06 mm**<br>**ns** |
| | No effect. | | No effect. | | No effect. | |
| 100- grain weight | 0.01 g + 0.02 g<br>= **0.03 g**<br>**ns + ns** | **0.26 g**<br>**ns** | 0.01 g + 0.17 g<br>= **0.18 g**<br>**ns + down** | **0.47 g**<br>**down/down** | 0.01 g + 0.02 g +0.17 g<br>= **0.21 g**<br>**ns + ns + down** | **0.28 g**<br>**down** |
| | No effect. | | There is an enhancing effect of *osxlg1* to *osxlg3b.* | | There is an enhancing effect of *osxlg1* and *osxlg3a* to *osxlg3b.* | |
| Yield | 5 g + 16 g<br>= **21 g**<br>**down + down** | **13.4 g**<br>**down** | 5 g + (-7.1 g)<br>= **-2.1 g**<br>**down + up** | **8.67 g**<br>**down** | 5 g+ 16 g + (-7.1 g)<br>= **13.5 g**<br>**down + down + up** | **14.37 g**<br>**down** |
| | *osxlg1* and *osxlg3a* are partially redundant. | | *osxlg1* is epistatic to *osxlg3b* with an enhancing effect of *osxlg3b* to *osxlg1.* | | *osxlg3b* is epistatic to *osxlg1* and *osxlg3a.* | |
| Biomass | (-23 g) + 71.4 g<br>= **48.1 g**<br>**ns + ns** | **14.5 g**<br>**ns** | (-23 g) + (-205 g)<br>= **-227 g**<br>**ns + up** | **-77.3 g**<br>**up** | (-23 g) + 71.4 g + (-205 g)<br>= **-156.5 g**<br>**ns + ns + up** | **-52.5 g**<br>**up** |
| | No effect. | | There is an enhancing effect of *osxlg1* to *osxlg3b.* | | *osxlg1* and *osxlg3a* suppress *osxlg3b.* | |
| Harvest index | 0.06 + 0.12<br>= **0.17**<br>**down + down** | **0.09**<br>**ns** | 0.06 + 0.15<br>= **0.21**<br>**down + down** | **0.15**<br>**down** | 0.06 + 0.12 + 0.15<br>= **0.33**<br>**down + down + down** | **0.19**<br>**down** |
| | *osxlg1* and *osxlg3a* are epistatic. | | *osxlg1* and *osxlg3b* are partially redundant. | | *osxlg1, osxlg3a* and *osxlg3b* are partially redundant. | |
